# Single cell RNA-sequencing reveals no evidence for meiotic sex chromosome inactivation in the threespine stickleback fish

**DOI:** 10.1101/2024.11.26.625488

**Authors:** Daniel E. Shaw, Wynter D. Ross, Alexis V. Lambert, Michael A. White

**Affiliations:** Department of Genetics, University of Georgia, Athens, GA, 30602

## Abstract

Sex chromosomes often evolve unique patterns of gene expression during spermatogenesis. In many species, sex-linked genes are downregulated during meiosis in response to asynapsis of the heterogametic sex chromosome pair (meiotic sex chromosome inactivation; MSCI). This process has evolved convergently across many taxa with independently derived sex chromosomes. Our understanding how quickly MSCI can evolve and whether it is connected to the degree of sequence degeneration remains limited. Teleost fish are a noteworthy group to investigate MSCI because sex chromosomes have evolved repeatedly across species, often over short evolutionary timescales. Here, we investigate whether MSCI occurs in the threespine stickleback fish (*Gasterosteus aculeatus*), which have an X and Y chromosome that evolved less than 26 million years ago. Using single-cell RNA-seq, we found that the X and Y chromosomes do not have a signature of MSCI, maintaining gene expression across meiosis. Using immunofluorescence, we also show the threespine stickleback do not form a condensed sex body around the X and Y, a feature of MSCI in many species. We did not see patterns of gene content evolution documented in other species with MSCI. Y-linked ampliconic gene families were expressed across multiple stages of spermatogenesis, rather than being restricted to post-meiotic stages, like in mammals. Our work shows MSCI does not occur in the threespine stickleback fish and has not shaped the evolution of the Y chromosome. In addition, the absence of MSCI in the threespine stickleback suggests this process may not be a conserved feature of teleost fish, despite overall sequence degeneration and structural evolution of the Y chromosome, and argues for additional investigation in other species. We also observed testis-dependent differences in coding and expression evolution for X-linked genes, revealing evidence of testis specific faster-X effect and gene-by-gene dosage compensation.

**Author Summary:** As male germ cells enter meiosis, the X and Y chromosome of many species undergo a drastic repression of gene expression. In mammals, this process has been shown to be essential for fertility, and the expression of sex-linked genes can lead to meiotic arrest and cell death. This process has only been studied in a handful of organisms, which limits our understanding how extensive MSCI is across the tree of life. Teleost fish are an understudied group with many examples of independently derived sex chromosomes across closely related species. Here, we investigate whether MSCI occurs in the threespine stickleback fish, using single-cell transcriptional profiling. We found gene expression remains active throughout meiosis on the sex chromosomes, indicating MSCI does not occur. This indicates that MSCI is not a conserved feature of all taxa and is not an inevitable outcome of degenerating Y chromosomes.

## Introduction

Heteromorphic sex chromosomes often fail to fully synapse during meiosis due to a lack of sequence homology between the chromosome pair (i.e. X and Y or Z and W). In mammals, the X and Y chromosomes only synapse within the pseudoautosomal region, where all crossing over is restricted in males [1]. Outside of the pseudoautosomal region, the unsynapsed chromatin triggers a process of complete transcriptional silencing and sequestration into a dense sex body, known as meiotic sex chromosome inactivation (MSCI) [2–4]. Most genes remain silenced throughout post-meiotic stages [5, 6] MSCI may actually represent a more generalized process of silencing of any unsynapsed chromosomes in male or female meiosis (MSUC; meiotic silencing of unsynapsed chromatin) [7, 8]. Although the function of silencing remains unknown, it may have a role in ensuring DNA double strand breaks are properly repaired during meiosis at locations where chromosomes fail to synapse properly (reviewed in [3].

MSCI appears to have evolved independently in multiple lineages. In mammals, MSCI evolved in the common ancestor of therians, approximately 180 million years ago after the origin of the X and Y chromosomes [9]. Since their origin, the sex chromosomes have diverged structurally in both marsupial and eutherian lineages, yet they still share many meiotic silencing factors that initiate in response to the unpaired X and Y [10, 11]. Sex chromosomes independently evolved within monotremes, and while a canonical sex body does not form [12], gene expression is reduced drastically in the non-recombining region in platypus [13]. This indicates a general mechanism for silencing unpaired chromosomes may have been present in the common ancestor of all mammals. Outside of mammals, sex chromosome silencing occurs across the hemizygous X chromosomes of *Caenorhabditis* species [14] and grasshoppers [15, 16], both of which have XO males, where the X lacks a pairing partner. Some evidence exists for transcript repression from the X chromosome in the XY male germline of *Drosophila melanogaster* [17–20], suggesting MSCI may also exist in this species (but also see [21, 22]).

MSCI imposes a strong selection pressure on the overall gene content of evolving sex chromosomes. The X chromosome is a disfavored environment for genes with functions in spermatogenesis if the X is transcriptionally silenced during meiosis [3]. One way to circumvent silencing is to move genes from the suppressed sex chromosomes to the autosomes.

Retrotransposition of genes has been a major driver of gene content evolution on the mammalian and *Drosophila* X chromosomes [23–25]. Another mechanism to circumvent transcriptional suppression during meiosis may be to increase copy number of genes on the X chromosome to produce more transcripts in spite of the repressive chromatin environment [3]. Many multi-copy gene families on the mouse X chromosome are reactivated post-meiotically despite chromosome-wide transcriptional silencing [26, 27]. Large multi-copy gene families also exist on the mouse and bull Y chromosomes that have testis-specific gene expression and have roles in spermatogenesis [28, 29]. Sex chromosomes have evolved repeatedly throughout the tree of life, but outside of a few well-characterized species with relatively ancient sex chromosomes it remains unclear the overall extent to which MSCI occurs and whether it is a common selection pressure on sex-linked gene evolution.

The degree of synapsis between sex chromosomes can even vary in species where substantial degeneration has occurred on the Y or W chromosomes. Birds have a ZW sex chromosome system that evolved independently from mammals over 120 million years ago [30]]. Similar to the mammalian Y chromosome, many species of bird W chromosomes share little sequence homology with the Z chromosome [30]. Despite this, MSCI does not occur on the ZW sex chromosomes [31]. In chickens, the heterologous Z and W chromosomes show complete synapsis in a majority of pachytene oocytes [31]. Incomplete MSCI has also been shown within mammals, where self-synapsis of the X chromosome in some spermatocytes blocks transcriptional suppression of the chromosome [32]. Sex chromosomes with little sequence divergence may also synapse more readily during meiosis, preventing MSCI. At the most extreme, sex chromosomes that only differ at a small sex determination region may not interfere with synapsis at all. For example, there is little sequence divergence between the X and Y chromosome in the guppy [33–35], *Poecilia reticulata*, and these chromosomes fully pair with each other during meiosis [36]. Additional focus on how different sex chromosomes achieve pairing during meiosis is necessary to understand the extent of MSCI across convergently evolving sex chromosomes.

The threespine stickleback fish *(Gasterosteus aculeatus)* is an excellent species to explore whether MSCI occurs on a degenerating sex chromosome that fully synapses. The threespine stickleback has an X and Y sex chromosome that evolved less than 26 million years ago [37]. Since this time, the Y chromosome has diverged from the X chromosome at both the sequence and structural level. The X and Y chromosomes have a 2.5 Mb pseudoautosomal region, where all crossing over is restricted during meiosis [38]. The non-crossover region on the Y chromosome has lost nearly half of the ancestral gene content [37, 39]. In addition, three pericentric inversions have altered synteny between the X and Y [37, 40]. Despite this degeneration, the sex chromosomes fully pair in a non-homologous fashion along their entire chromosomal length during prophase I of meiosis [41]. Threespine stickleback provides a needed perspective on the evolution of MSCI within teleost fish. Due to sex chromosome pairing in meiosis, we hypothesized that MSCI would not occur during spermatogenesis and consequently this process has not shaped the evolution of gene content on these chromosomes.

In this study, we used single-cell RNA sequencing to characterize sex-linked gene expression throughout spermatogenesis. Consistent with full pairing and synapsis, we did not find a stage-specific down regulation of X- or Y-linked genes across spermatogenesis. The sex chromosomes were also not sequestered into a sex body. Our study indicates asynapsis and MSCI is not an inevitable outcome for all degenerating sex chromosomes and argues for additional research into other sex chromosome systems to understand why some heteromorphic sex chromosomes can maintain synapsis despite a lack of sequence homology.

## Results

### Identification of stickleback spermatogenic stages using single-cell RNA sequencing

The testis is a highly heterogeneous organ and gene expression varies both spatially across cell types as well as temporally across developmental stages [13, 42]. Spermatogenesis occurs synchronously in the threespine stickleback [43]. At a given developmental stage, the testes will be enriched for premeiotic, meiotic, or post-meiotic cells, along with somatic support cells. In reproductively mature adult males, the testes will mostly contain mature sperm and somatic cells. In order to capture all stages of spermatogenesis, we sequenced testes from lab-reared males at different developmental stages. We sequenced 6147 cells from a single testis dissected from an earlier stage of spermatogenesis and 2652 cells from a single testis dissected from a male at a later stage of spermatogenesis. In order to maximize the total number of stages of spermatogenesis, we also pooled the testes from two additional males from an intermediate developmental stage (6027 cells total). The median number of genes detected were comparable across samples (Testis sample 1: 271, Testis sample 2: 311, Testis sample 3: = 281), but sample two had a higher number of genes and UMIs detected (Supplemental Figure 1), likely due to this being the sample with lowest number of cells. Each testis sample cannot be treated as a direct replicate of one another due to the differences in timing of synchronized spermatogenesis and variation in the number of cells. However, we did find many replicated clusters, indicating overlap in stages of spermatogenesis between the testis samples (Supplemental Figure 2).

Consistent with this, cells were hierarchically clustered by type rather than by sample, when considering gene expression (Supplemental Figure 3). This suggests that batch effects on expression level were minimal after integration and normalization. While we found similar cell types across each sample, we found variation in the density of each cell type across samples. As expected, we found a higher number of meiotic cells relative to post-meiotic cells, in the earliest developmental sample (sample 1). However, some differences in cell population may also be due to technical differences in sequencing library and UMI counts, as we also found less immune cells in sample 1, which also had the lowest amount of UMIs detected. Among the three samples, there were two unique cell clusters. Because these clusters were only present in a single sample, we removed the cells (cluster 9 from sample 1, cluster 6 from sample 3) from all downstream analyses. Non-replicated clusters could be due to rare cell populations only identified in one sample or derived from batch differences between sequencing libraries [44].

To capture all stages of spermatogenesis in a single analysis, we merged the three samples together and scaled gene expression accordingly. Between all of the samples, we recovered a total of 12,136 cells after filtering. Clustering analysis revealed 15 distinct cell clusters (Figure 1A). We identified 3,694 differentially expressed genes across all cell clusters (Mann-Whitney U Test) (Supplemental File 1). Genes were considered differentially expressed within a cluster if the gene was expressed in 25% of cells within the cluster, had a log_2_ fold change > 0.25 for expression compared to all other clusters, and had a significantly different level of expression evaluated with a P-value < 0.01 after correcting for multiple comparisons. Among the differentially expressed genes, we identified those unique to cell-types in order to assign spermatogenesis stages to each cluster.

**Figure 1.**
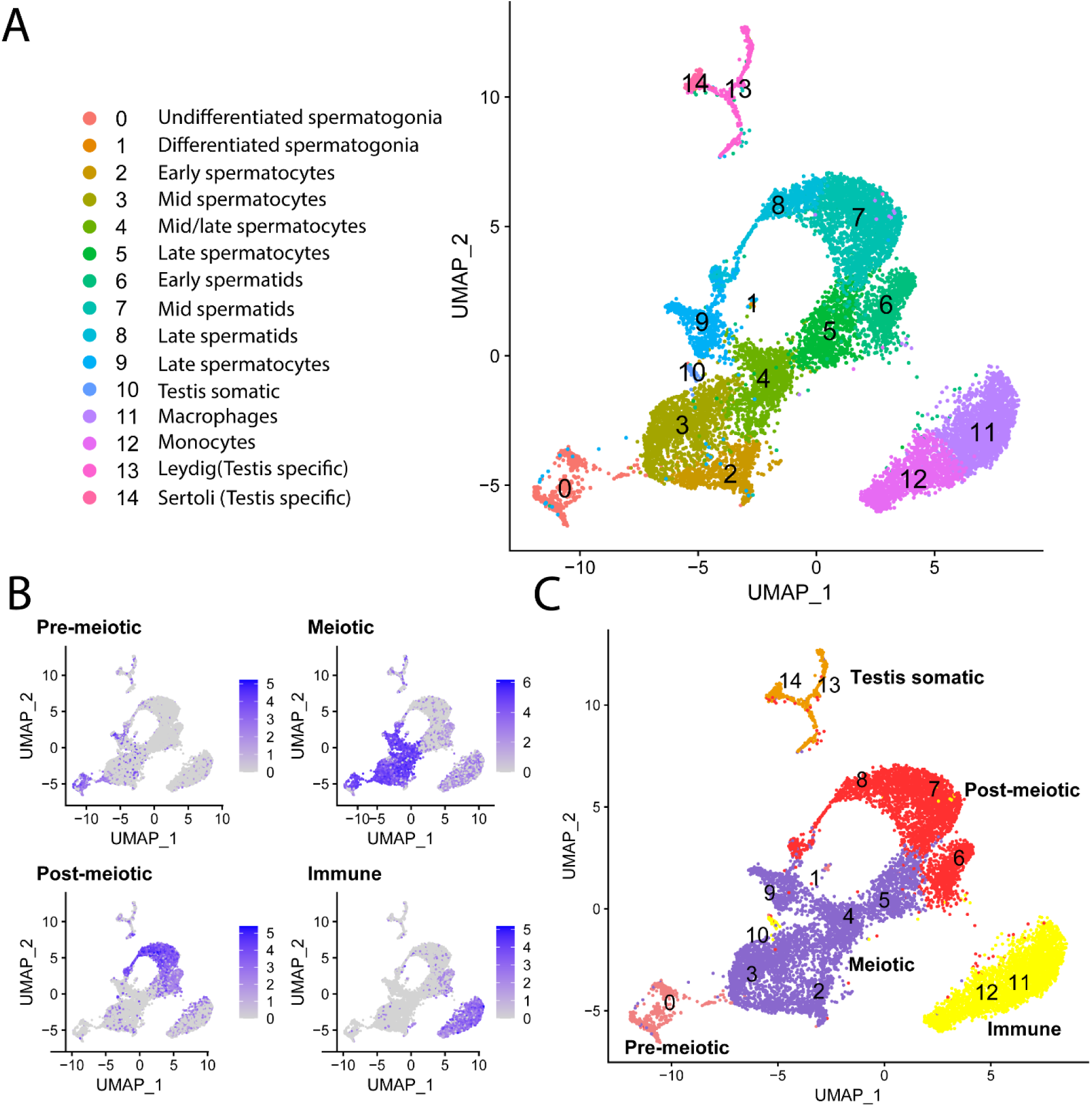
Single-cell RNA-seq of threespine stickleback spermatogenesis. Identified clusters of pre-meiotic, meiotic, post-meiotic, and somatic cell populations integrated from three sequencing runs. (**A**) Visualization of single-cell expression profiles of spermatogenesis in UMAP projection. We identified 15 clusters and related them to stages of spermatogenesis based on significantly upregulated marker genes and GO terms (Table 1). (**B**) Examples of enrichment of key marker genes across the four main stages of spermatogenesis. Additional marker genes are in Table 1. (**C**) The four main spermatogenesis stages overlaid on the UMAP projection.

**Table 1.**
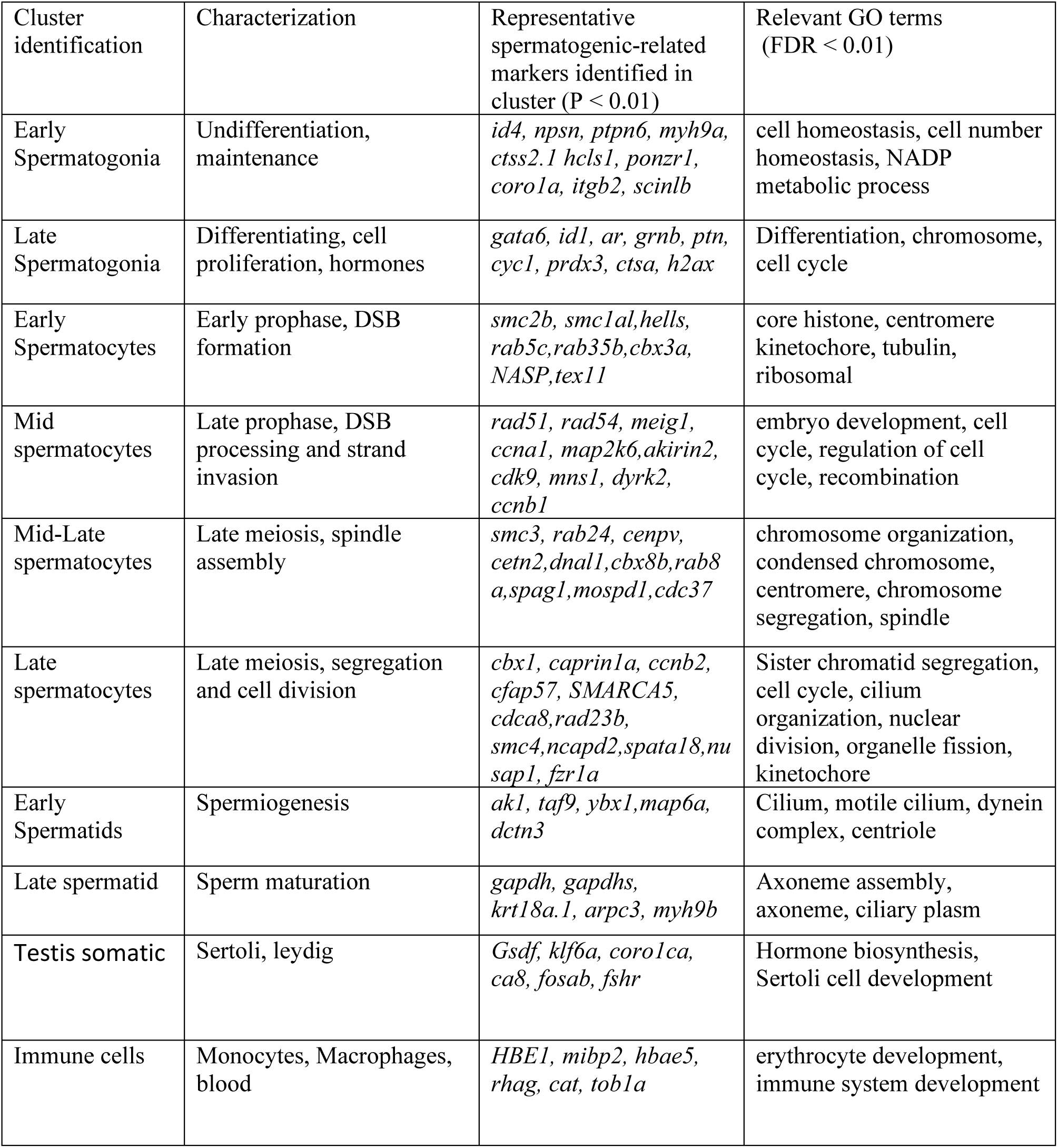
Representative marker genes and gene ontology functions enriched in each cell type.

Cell-type specific markers were identified from conserved genes expressed during spermatogenesis from insects [45, 46], mammals [13, 42, 47–50], and other teleost fish [51–56]. Using these markers, we identified spermatogonia, spermatocytes, spermatids, and somatic cell types (Table 1; Supplemental Figure 4). One class of somatic cells clustered separately from spermatogenesis clusters and had differentially expressed genes that were not specific to testis (i.e. *Hbae5*, *Rhag*; Cluster 11 and 12; Figure 1A). These were genes that are mostly expressed in immune cells (e.g. monocytes and macrophages). We confirmed the expression of these markers during spermatogenesis in other teleost species, including *Sebastes melanops* (black rockfish) [55] and *Danio rerio* (zebrafish) [51]. These genes were also expressed in cells of the head kidney, a hematopoietic organ with immune function in threespine stickleback [56].

A second group of somatic cells mapped distinctly from spermatogenesis cells, but also had some expressed genes that were shared with some of the spermatogenesis clusters, consistent with somatic cells of the testis (Sertoli and Leydig cells; Cluster 13; Figure 1A). The markers we used to identify Sertoli and Leydig cells (e.g. *Gsdf* and *Fshr)* were also confirmed to be expressed in zebrafish [51] and rockfish [55]. Within the differentially expressed genes, we found several significantly enriched functions that corroborated our cell type assignment (Table 1).

For further analyses, we combined clusters into four major cell groups to explore differences across major cell types: pre-meiotic, post-meiotic, meiotic, and somatic cells (Figure 1B and 1C). Within each of these broader categories, we found that gene expression levels were highly correlated across each testis sample before integration (Spearman Rank Correlation R > 0.65 for all comparisons) (Supplemental Figure 5). We found the major cell type groups were concordant between samples. The average gene expression of the 2000 highest expressed genes in major cell types was highly correlated after normalization (mean rho of shared clusters = 0.83, all vs all clusters average rho = 0.36, Supplemental file 2).

### Sex-linked genes do not undergo meiotic silencing

In mammals, genes within the non-recombining region of the sex chromosomes are not transcribed (MSCI) [2, 3, 8]. To determine whether meiotic silencing has also evolved in threespine stickleback fish, we compared rates of transcription between sex-linked and autosomal genes for all genes expressed in that cell type. We found that gene expression was not reduced on the X chromosome in meiotic cells, relative to somatic and pre-meiotic cell types (Figure 2; P > 0.05). While the Y chromosome showed lower expression than the X chromosome, Y-linked genes maintain similar levels of expression across pre-meiotic and meiotic cells (Figure 2, P > 0.05). We also investigated whether MSCI was operating at a finer scale within cell types or if this pattern was restricted to the four major cell groups. While we did detect more variation in X:AA expression between cell types when analyzing each sample, likely due to the smaller sample size of each cluster, we found that meiotic and post-meiotic expression was not significantly lower than pre-meiotic expression in all three samples (Supplemental Figure 6), indicating MSCI was also not operating in individual cell types.

**Figure 2.**
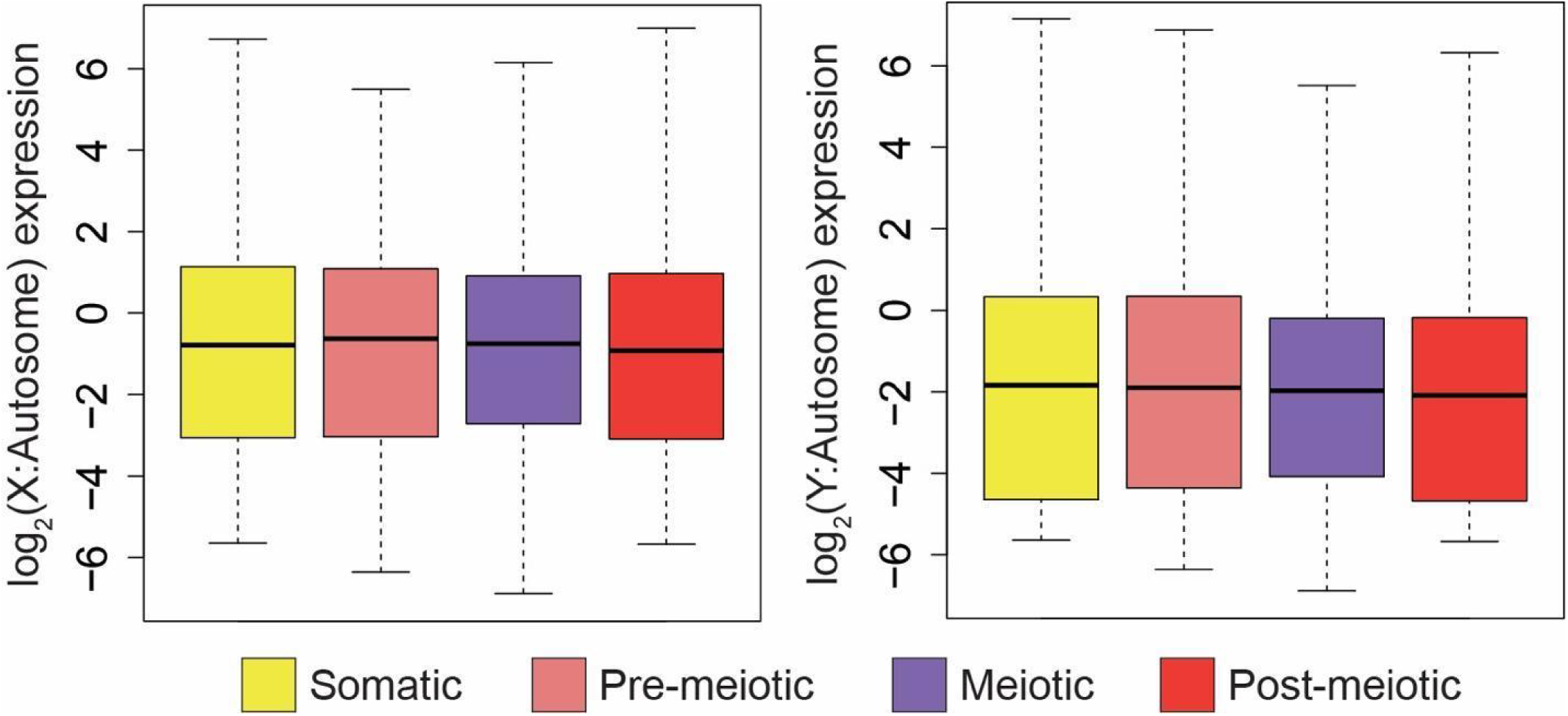
Sex-linked gene expression does not change throughout spermatogenesis. X- and Y-linked gene expression was normalized to median autosome expression level for each stage of spermatogenesis. Relative to median autosome expression, X- (left) and Y-linked (right) gene expression does not change across stages of spermatogenesis. (Pairwise Mann-Whitney U Test, P > 0.05). Outliers are not shown. Pre-meiotic X: N = 500, Meiotic X: N = 506, Post-meiotic X: N = 534, Somatic X: = 524. Pre-meiotic Y: N= 442, Meiotic Y: N = 468, Post-meiotic Y: N = 442, Somatic Y: N = 410. Analysis excludes clusters 6 and 9 due to lack of replication between samples.

MSCI could be restricted to certain classes of genes or regions of the sex chromosomes. For example, testis-specific genes could be expressed higher, while housekeeping genes are repressed. To test for this, we filtered out genes that appear to be exclusively expressed in testis bulk RNA-seq compared to brain and liver [37, 39]. We found that broadly expressed genes are also expressed across meiosis (Supplemental Figure 7).We also examined whether there were unique expression patterns among the differently aged evolutionary strata (Figure 3) [37]. We found no evidence of silencing among the different regions of the sex chromosomes (Figure 3), indicating this is not a phenomenon restricted to even the oldest stratum of the X and Y (stratum 1). Next, we repeated the analysis using a distinct clustering algorithm within Seurat, and by clustering with a different software package (scanpy [57]). In both cases, we found that normalized X expression was similar across cell types of spermatogenesis, regardless of cell clustering approach and cell type annotation (Supplemental Figure 8). Overall, we found that unlike mammals, chromosome-wide silencing does not occur on the threespine stickleback sex chromosomes during meiosis.

**Figure 3.**
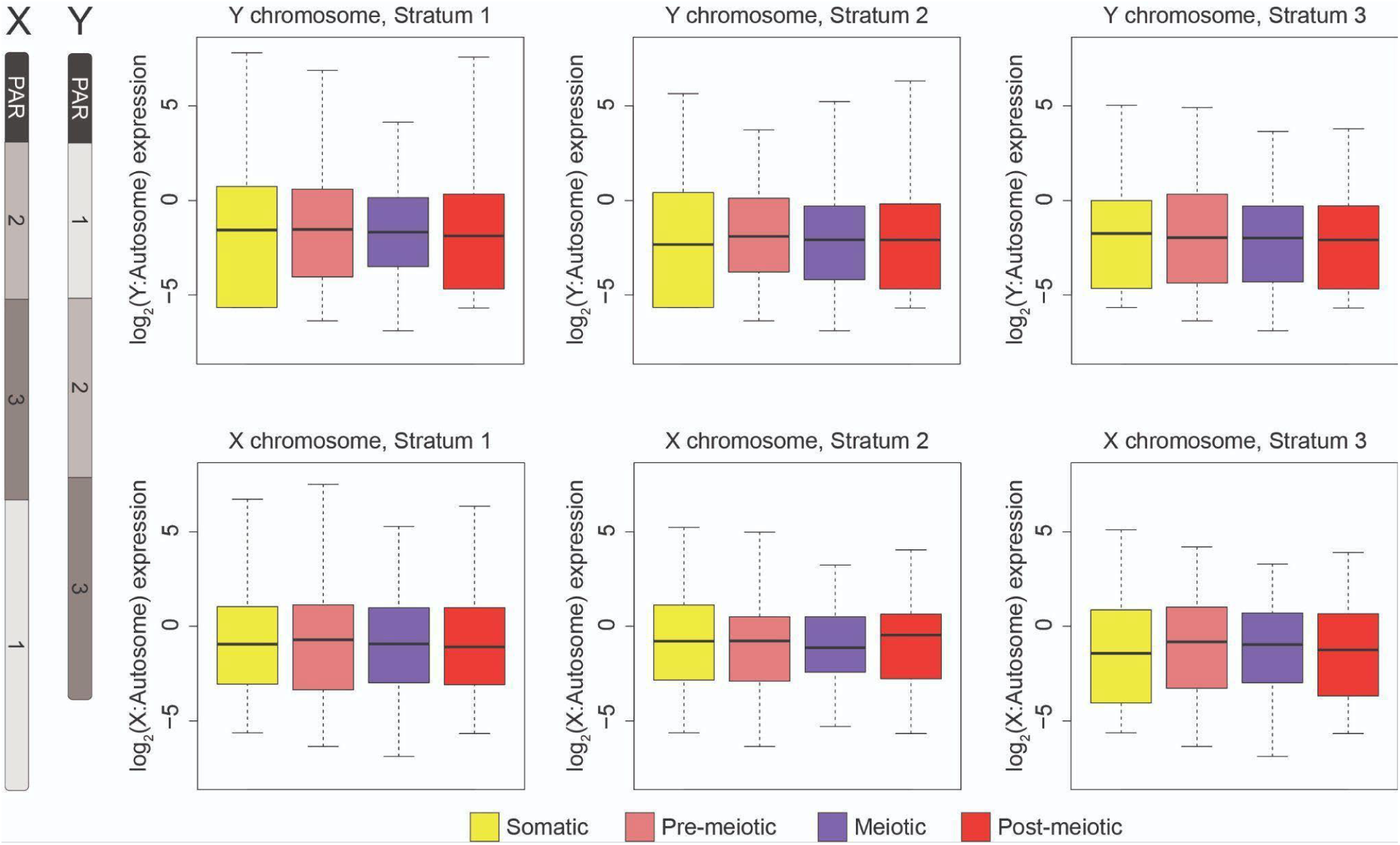
Sex-linked gene expression does not change throughout spermatogenesis in each of the evolutionary strata. The sex chromosomes contain three distinct evolutionary strata at different states of divergence formed by nested inversions (left). Sex-linked gene expression does not change across spermatogenesis in the three different evolutionary strata. X- and Y-linked gene expression was normalized to median autosome expression level for each stage of spermatogenesis. Relative to median autosome expression, X- (bottom) and Y-linked (top) gene expression does not change throughout spermatogenesis in each of the three strata (stratum 1: oldest, stratum 2: middle, stratum 3: youngest; Pairwise Mann-Whitney U Test, P > 0.05). The recombining pseudoautosomal region (PAR) is indicated for reference. Outliers are not shown.

### The threespine stickleback X and Y chromosomes are not condensed into a sex body

During MSCI, the mammalian X and Y chromosomes are sequestered into a dense structure, known as the sex body [2–4, 8]. A key marker of this structure are the phosphorylated histone variant, γH2AX [2, 3, 6, 58]]. The threespine stickleback X and Y chromosomes likely synapse fully and do not show evidence of delayed DSB repair [41]. Together with our results that gene expression was not suppressed throughout meiosis, we predicted that a mammalian-like MSCI sex body would also not be present during spermatogenesis. To test this, we characterized how γH2AX accumulated throughout meiosis using immunofluorescence. γH2AX broadly localized to all chromosomes beginning at leptotene in response to the initiation of DNA double strand breaks [59] (Figure 4). During pachytene, double strand breaks are repaired and we observed a concurrent reduction of γH2AX foci, similar to patterns observed in mammals [2].

**Figure 4.**
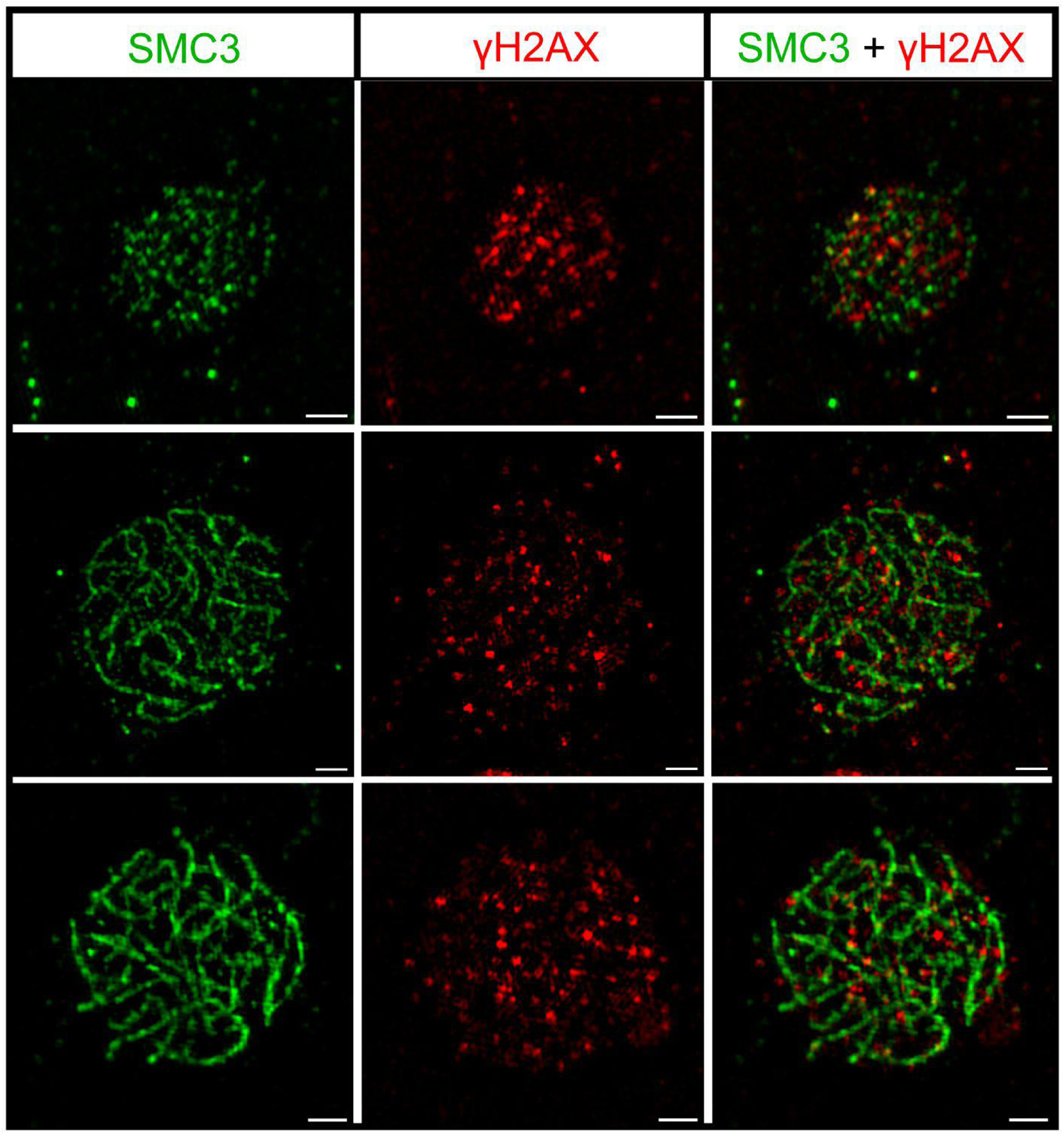
The X and Y chromosomes are not sequestered into a sex body. The highest density of the phosphorylated histone variant, γH2AX, a marker of DNA damage (red) occurs during leptotene (top), when the greatest number of DNA double strand breaks also occur. γH2AX localization decreases as double strand breaks are repaired in zygotene (middle) and pachytene (bottom). The sex chromosomes are repaired at the same rate since they are not sequestered into a dense, γH2AX containing sex body at pachytene. The synaptonemal complex axes are immunolabeled against the axial protein, SMC3 (green). The scale bar is 5 µm.

However, at this stage we did not observe an aggregation of γH2AX synonymous with a sex body (Figure 4). We staged a total of 146 nuclei from leptotene to pachytene. While, the X and Y are not distinguishable at this resolution, none of the scored nuclei had any structure resembling a sex body. This result is consistent with the expression data, supporting that canonical MSCI does not occur in threespine stickleback fish due to the lack of asynapsis and sex body formation.

### Gene evolution on the sex chromosomes has not been shaped by MSCI

Unique expression patterns can evolve rapidly on sex chromosomes [39, 60–64]. Sex chromosomes can differentially accumulate pre- and post-meiotic genes [27, 65], gametologs can gain testis-specific expression [66, 67], and Y-linked gametologs rapidly degenerate, losing expression [68]. To explore this, we looked for enrichment of ancestral gametologs that were only expressed from either the X or Y chromosome at different stages of spermatogenesis. We found that pre- and post- meiotic cell types were over-enriched for Y-specific genes, while meiotic cell types were under-enriched (Pre-meiotic: 2x, P < 0.001; Post-meiotic: 1.24x, P = 0.003; Meiotic: -3.15x, P < 0.001; Somatic: 1x, P = 0.5; hypergeometric test) (Supplemental Figure 9). We observed an over-enrichment of X-specific gametologs in pre-meiotic cell types, but no other cell types were over- or under-enriched (pre-meiotic 1.39x, P = 0.005; meiotic: 1.16x, P = 0.084; post-meiotic: -1.13x, P = 0.11; somatic: 1.1x, P = 0.290; hypergeometric test) (Supplemental Figure 9). These results suggest early and late stages of spermatogenesis are prone to the masculinization of gene expression, but not meiotic stages.

In addition to the ancestral complement of gametologs shared between the X and Y, the Y chromosome has also acquired unique genes that originated from the autosomes [37]. After translocation to the Y chromosome, many of these genes have further duplicated and evolved testis-biased expression, suggesting a gain of spermatogenesis function [37]. In mammals, the ancient Y chromosome has also accumulated species-specific gene duplicates that are enriched for post-meiotic expression and have been speculated to be involved in intra-genomic conflicts between X and Y bearing sperm during spermatogenesis [26, 28, 29, 69–71]. Due to the absence of meiotic silencing in stickleback fish, we characterized how the expression of these Y-linked duplicates compared to mammalian amplicons. We looked at the overall expression of all paralogous copies within each gene family and assayed whether there were differences across the four major cell types of spermatogenesis. We found that the Y-linked duplicated genes were expressed across multiple cell types of spermatogenesis (Figure 5). Interestingly, in the absence of chromosome wide silencing in meiosis, we found the highest number of expressed duplicate families (104) in meiotic cell types, and expression for many duplicate families is highest in meiosis compared to all other cell types (Figure 5). This was true for many genes, regardless of the total number of paralogs, which ranged from 2-30 copies. This suggests that many amplicons on the threespine stickleback Y chromosome may have functions throughout meiosis, rather than post-meiotic stages as has been observed in mammals.

**Figure 5.**
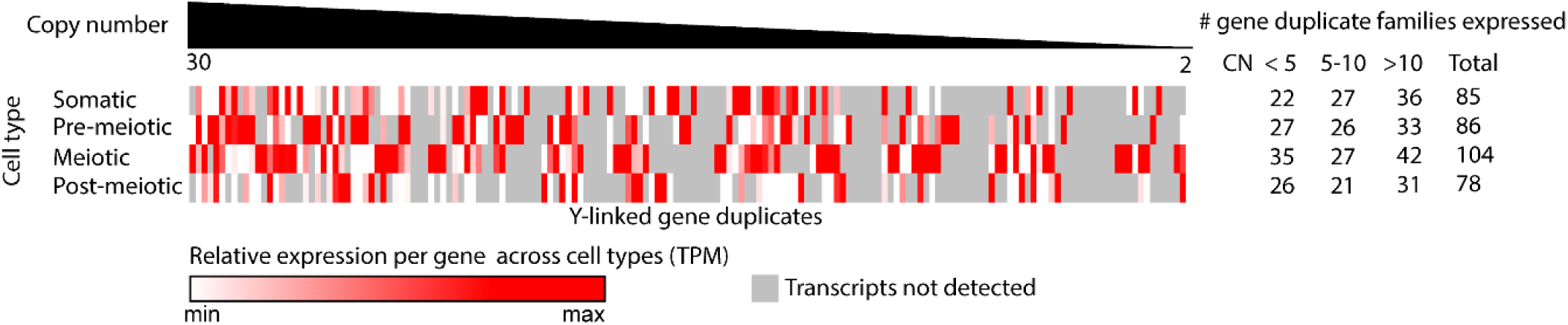
Y-linked duplicate gene families are expressed across stages of spermatogenesis. Duplicate gene families on the Y chromosome are enriched for expression during spermatogenesis. The greatest number of duplicate families were found to be expressed in meiotic cell types. Normalized counts for all paralogous copies of a gene family were summed to determine an aggregate expression level.

We searched for an enrichment of GO terms among the expressed duplicate genes to assign function. We found that the predominant functions included chromatin assembly, DNA packaging, and DNA repair. Specifically, we found many expressed duplicate families were genes with critical functions in meiosis, including proteins involved in double strand break repair (RAD proteins) and proteins involved in chromosome synapsis and segregation (SMC proteins) [72]. Together, these results provide novel insight into the functions of duplicated Y-linked genes, which in other species have been associated with post-meiotic expression [69, 70, 73]. Specifically, this finding highlights the role of meiotic functions in duplicated gene families expressed on the Y, in the absence of meiotic silencing.

We searched for evidence of retrotransposition of genes throughout the genome, which occurs at an increased frequency from the X chromosome in species with MSCI [17, 23, 74, 75]. Unlike these species, we did not find an excess of retrotransposition from the X chromosome in the threespine stickleback. We identified nine examples of retrotransposition since the threespine stickleback shared a common ancestor with the ninespine stickleback *(Pungitius pungitius)* approximately 26 million years ago, before the threespine sex chromosomes evolved (the ninespine stickleback has independently derived sex chromosomes) [76–78]. None of the nine retrotransposition events were from the X chromosome. While nine total retrotransposition events provides a low sample size to test for under-enrichment on the X chromosome, this trend is much less pronounced than in humans and mice [73] Even though the threespine stickleback Y chromosome shares many features of gene content evolution that have been observed on the mammalian Y chromosome [37, 39], our results highlight that MSCI has not been a driving force shaping these patterns.

### Genes expressed during spermatogenesis show signs of dosage compensation

Many species have evolved mechanisms to compensate for the loss of expression from the Y-linked allele [79]. However, threespine stickleback do not have evidence of chromosome-wide mechanisms of dosage compensation in the heterogametic males, at least within brain tissue analyzed in bulk [39]. It is possible that gene-by-gene dosage compensation has evolved and it may be restricted to certain tissue or cell types. Indeed, the extent of chromosome-wide dosage compensation has been shown to be variable across tissues [80–83]. To look for dosage compensation during spermatogenesis, we compared normalized expression of X-linked genes that still had a Y-linked allele (X+Y gametologs) with those where the Y-linked allele has been lost (X-only gametologs). If dosage compensation is not occurring during spermatogenesis, we would expect X-only gametologs to have expression levels half of what is observed globally across autosomes. Instead, X-only gametologs had expression levels more similar to autosomes. These X-only gametologs were also expressed significantly higher than X+Y gametologs in all four major cell groups (Figure 6 ; P < 0.01). We compared this finding across the differentially aged evolutionary strata of the sex chromosomes. If dosage compensation is occurring, we would expect this effect would be strongest in the oldest evolutionary stratum (stratum 1), where the Y chromosome has been degenerating for the greatest amount of time. Consistent with this, we only detected a significant difference between X+Y gametologs and X-only gametologs for all cell types in the oldest stratum (Stratum 1; P < 0.01) (Figure 6). This result is consistent with previous findings that *cis*-regulatory elements in testis exhibited faster rates of sequence divergence and positive selection on the X chromosome [65]. Our findings provide evidence that sex-linked genes expressed during spermatogenesis may be dosage sensitive and have evolved mechanisms to compensate for dosage loss from the Y chromosome.

**Figure 6.**
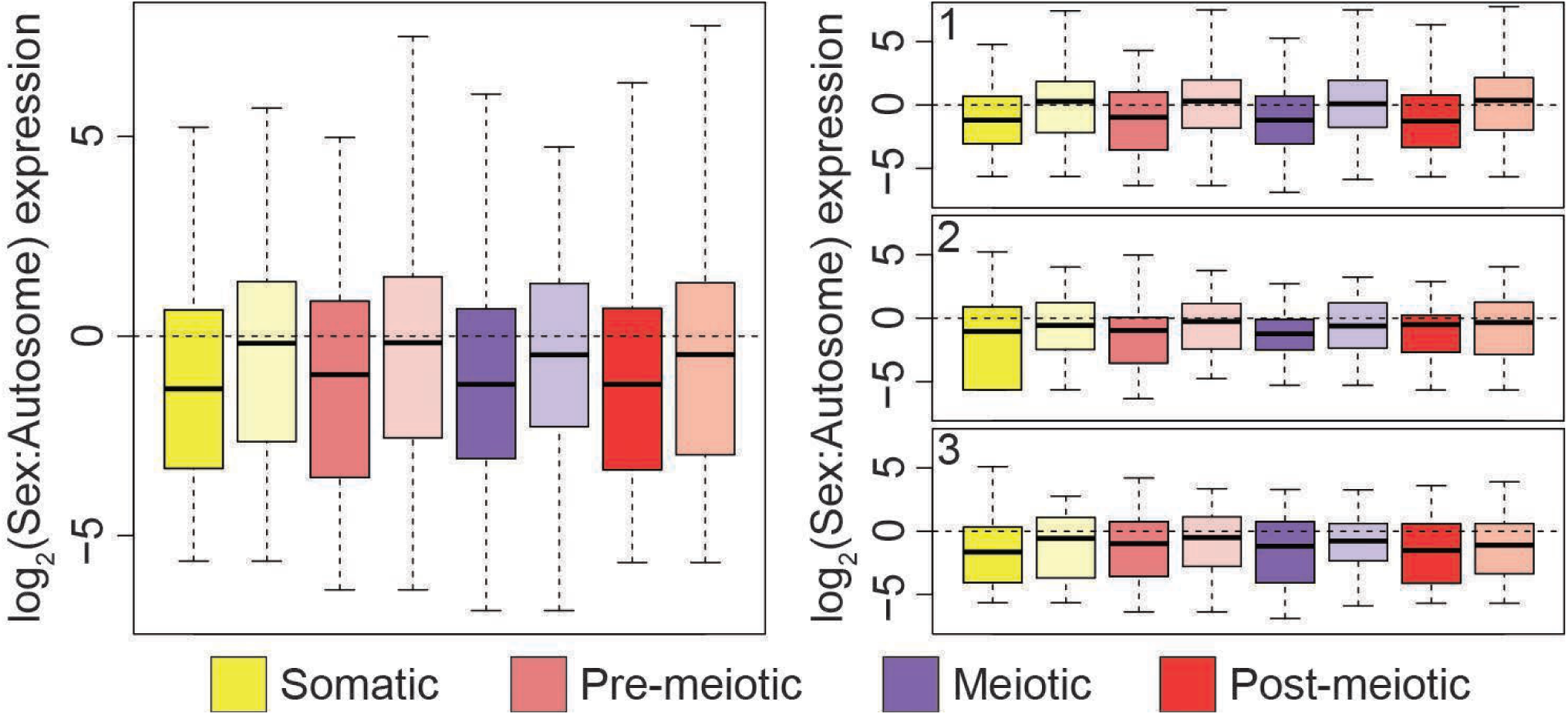
Spermatogenesis genes that have lost a Y-linked allele are dosage compensated. Genes with alleles on both the X and Y chromosomes (X+Y gametologs; dark shades) have lower expression relative to autosomes than genes that only have an X-linked allele (X-only gametologs; lighter shades). This indicates the X-only gametologs are upregulated to more closely match median autosome expression levels. All X-only genes are significantly higher than X+Y gametologs for all cell types (somatic: P = 0.004; pre-meiotic: P = 0.008; meiotic: P= 0.015; post-meiotic: P = 0.037; Kruskal-Wallis and Dunn’s test.). When separated by the three evolutionary strata (right panels), X-only genes are expressed higher for all cell types in stratum 1 (the oldest region). X-only genes and X+Y gametologs are not significantly different for strata 2 and 3 (the two youngest regions) across all cell types (stratum 1 somatic: P = 0.001; stratum 1 pre-meiotic: P = 0.007; stratum 1 meiotic: P= 0.012; stratum 1 post-meiotic: P = 0.004; stratum 2 somatic: P = 0.301; stratum 2 re-meiotic: P=0.124; stratum 2 meiotic: P = 0.31; stratum 2 post-meiotic: P = 0.391; stratum 3 somatic: P = 0.115; stratum 3 pre-meiotic: P = 0.34; stratum 3 meiotic: P = 0.302; stratum 3 post-meiotic: P = 0.632; Kruskal-Wallis and Dunn’s test). Outliers are not shown.

### Genes have similar signatures of selection throughout stages of spermatogenesis despite testis specific faster-X effect

Sex chromosomes that have recently evolved are less likely to have signatures of a faster-X effect [84], as XY shared genes are not hemizygous. Previous work in stickleback fish found some evidence for a faster-X effect among hemizygous genes in the youngest strata [39].

However, this study used variants discovered from short-read sequencing and was additionally limited by the lack of a reference threespine stickleback Y chromosome assembly and the lack of a ninespine stickleback whole-genome assembly as an outgroup. Additionally, sex-linked genes in the oldest part of the Y chromosome have been shown to be under purifying selection to maintain dosage-sensitive genes [37, 39]. With improved genome resources now available, we first wanted to test for evidence of different rates of coding evolution of sex-linked genes [37, 85, 86]. We identified all coding substitutions between threespine and ninespine stickleback, which has an independently evolved sex chromosome pair on chromosome 12 [76]. We found similar rates of non-synonymous to synonymous substitution rates (*d*_N_/*d*_S_) between autosomal genes and all X-linked genes that are hemizygous. Y-linked hemizygous genes had a higher *d*_N_/*d*_S_ than autosomes, consistent with the degeneration of coding regions on this chromosome (Supplemental Figure 10, P = 0.008). Interestingly, we found that X-linked gametologs that still maintain a Y copy had lower *d*_N_/*d*_S_ than autosomes (P < 0.001, Supplemental Figure 10). We also observed a lower *d*_N_/*d*_S_ than average for the X chromosome in the ninespine stickleback (chromosome 12) (Supplemental Figure 11), suggesting X-linked gametologs are evolving slower in both species. This is consistent with a recent theoretical model that proposed selection is less efficient for gametologs on the X chromosome [83] (i.e. slower-X effect). Interestingly, *d*_N_/*d*_S_ of X-linked hemizygous genes was higher than autosomes, only for genes induced in major cell groups of spermatogenesis, suggesting that the faster-X effect has affected male-specific genes (Figure 7, P = 0.018, Kruskal-wallis and Dunn’s test).

**Figure 7.**
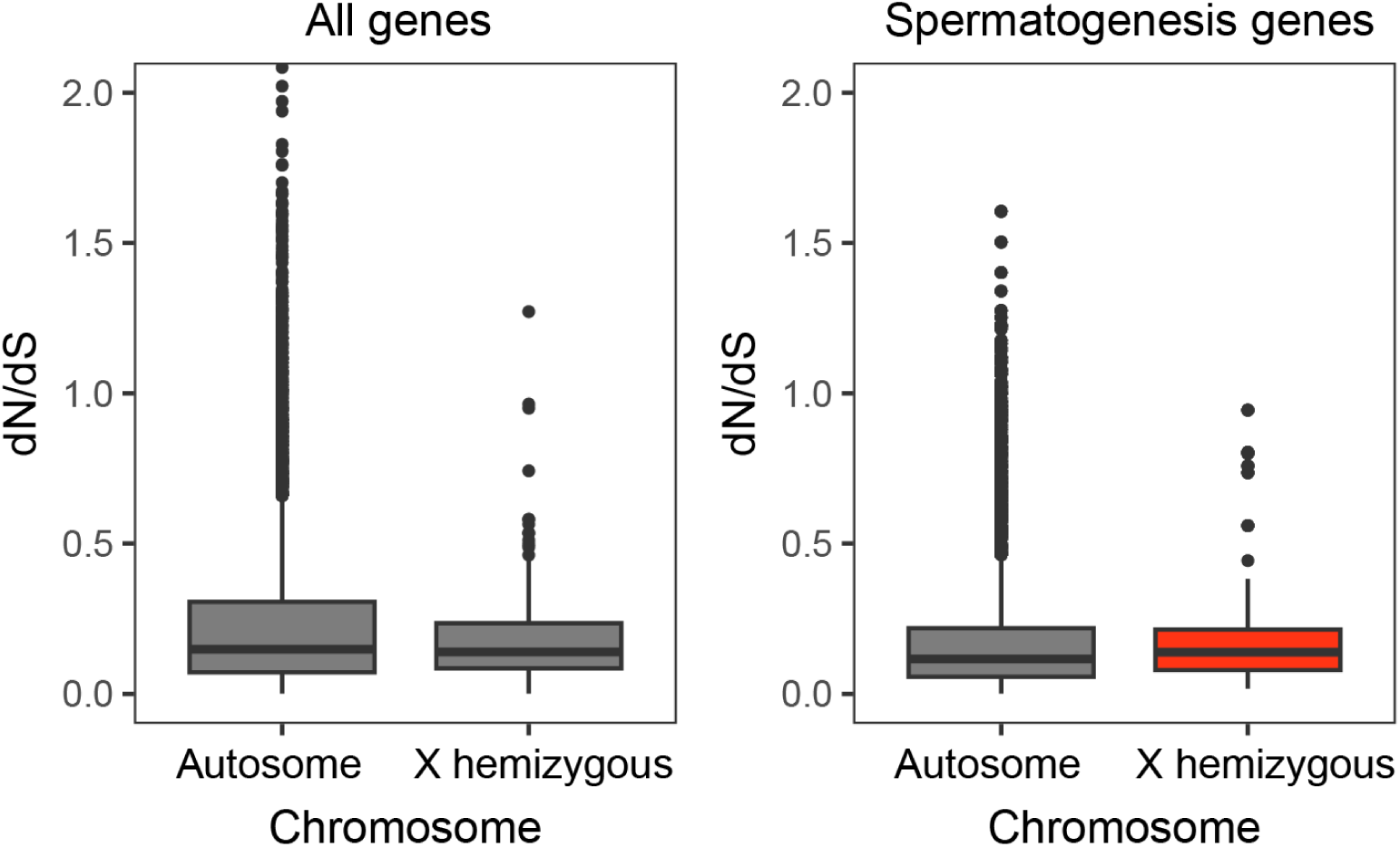
The faster-X effect is specific to spermatogenesis genes. Non-synonymous and synonymous substitutions were quantified between the threespine and ninespine stickleback fish, for 11,493 genes. X hemizygous genes have similar *d*_N_/*d*_S_ as autosomes for all genes. For genes induced in spermatogenesis (N = 2378), X hemizygous genes have higher *d*_N_/*d*_S_ than induced genes on autosomes (P = 0.018, Mann-Whitney U test, see Supplemental Figure 11 for all comparisons). Genes were defined as induced if they had expression levels greater than 2x the median expression in any major cell group in spermatogenesis.

In other species, stages of spermatogenesis have different rates of coding evolution [13, 87–89]. In house mice, cell types in early stages are more evolutionarily constrained, while later stages show signatures of rapid evolution, and this pattern is more extreme on the X chromosome due to the faster X effect of highly degenerated Y chromosomes [88]. However, it remains unclear if faster evolutionary rates would also apply to meiotic or post-meiotic genes on younger chromosomes in the absence of meiotic silencing. We tested for differences in evolutionary rates between genes induced across the four major cell groups of spermatogenesis. We found that *d*_N_/*d*_S_ was similar across all stages (Supplemental Figure 12, P = 0.501, Kruskal Wallis test).

Together, this suggests evolutionary rates are similar across stages of stickleback spermatogenesis.

## Discussion

The repression of sex chromosomes during meiosis was first noted based on early cytological evidence that many sex chromosomes were found in a condensed chromatin state [90–92]. Condensation of the X chromosome was proposed to be a critical feature of spermatogenesis, and could result in sterility if disrupted [90]. Since then, there have been many hypotheses how the function of MSCI may improve the fidelity of meiosis including to help avoid ectopic recombination, facilitate DSB repair, avoid expressing genes that have unrepaired DSBs, or act as a meiotic checkpoint response [3, 91]. Molecular studies have revealed that MSCI in mouse is initiated by the unsynaped state of the X and Y in male meiosis during the pachytene stage and results in a complete lack of gene expression [2, 8]. Other hypotheses for MSCI suggest it may be a result of how sex chromosome sequence evolves [93–95]. The silencing of the X and Y may limit the effect of segregation distorters that would bias the transmission of either chromosome during male meiosis [93]. The loss of spermatogenesis genes on the X chromosome, perhaps through sexually antagonistic selection, may favor the deactivation of the X during meiosis [94].

In this study, we found that the threespine stickleback X and Y chromosome maintain relative expression of genes across all stages of spermatogenesis. Our findings align with previous observations that the X and Y chromosomes fully pair during meiosis [41]. If the sex chromosomes are fully associating in a non-homologous fashion, this would prevent MSCI from occurring. In other species with heteromorphic sex chromosomes, full pairing has been shown to occur despite an overall lack of homology between the chromosomes. Each documented case thus far has revealed different mechanisms to achieve this. In the platypus, for instance, cohesins (SMC3) differentially accumulate on the five X and five Y chromosomes during meiosis [96].

Although the exact function of the increased recruitment of cohesins to the sex chromosome axial cores remains unknown [12, 96]. The threespine stickleback sex chromosomes do not exhibit differential accumulation of SMC3 between the sex chromosomes and autosomes [41], suggesting this is not required in all species for non-homologous pairing. In the chicken, the Z chromosome undergoes synaptic adjustment to pair with the W [92]. This is followed by assembly of the central element of the synaptonemal complex, indicating full synapsis of non-homologous regions of the chromosomes [31, 97]. Consequently, MSCI does not occur [31]. Synaptic adjustment of the longer sex chromosome is also observed during pairing in the threespine stickleback fish [41]. However, it remains unknown if full synapsis is also achieved between the X and Y [41]. Both the X and Y chromosomes appear closely associated and synapsed during prophase I of meiosis based on localization of the axial element, SMC3, but without higher resolution microscopy, it cannot be ruled out that some regions are not fully synapsed [41]. Additional work is necessary to understand if central element components, like SYCP1, are also recruited during non-homologous pairing of the threespine stickleback X and Y chromosomes and if full synapsis is required to prevent MSCI from occurring.

Alternatively, it is possible MSCI is not a conserved feature of all taxa and it does not occur in stickleback fish. The initial finding that MSUC occurred in taxa as diverse as *Neurospera crassa, C. elegans*, and mammals lead researchers to believe that MSCI may be initiated in all species with divergent, heteromorphic sex chromosomes [98]. However, evidence supporting the evolution of MSCI outside of the mammalian lineage has been somewhat confounding. Despite some evidence of lower expression [97], the W and Z chromosome in chicken oogenesis was found to largely maintain expression as the sex chromosomes maintain non-homologous synapsis in female oocytes [31], perhaps through a similar mechanism of synapsis as stickleback fish. In *Drosophila melanogaster*, some evidence suggests gene expression is lowered on the X chromosome during meiosis [18, 99–101]. However, the extent of silencing is less pronounced than mammals, does not occur on the Y, and may be an artifact of the lack of dosage compensation in spermatocytes, or the movement of spermatocyte genes off the X chromosome to avoid the lack of dosage compensation [22, 82, 83, 100]. Similar to our findings in stickleback fish, the neo-sex chromosomes in *D*. *miranda* show no evidence of MSCI [82]. Multiple insects have been reported to have silencing, including the grasshopper (*Eyprepocnemis plorans*), which has complete silencing, independent of γH2AX accumulation [102], and the mosquito (*Anopheles gambiae)* which has lower expression of the X chromosome, though not all genes are silenced [103, 104]. This highlights a need for additional species to be studied to understand the generalities of sex-linked gene expression during meiosis.

Teleost fish are one such understudied group. The existence of MSUC and MSCI remains unclear and has only been investigated in a few species [105]. The swamp guppy (*Micropoecilia picta*) has a degenerated Y chromosome relative to the X chromosome, yet cytological investigations of meiosis revealed no presence of an obvious sex body that would indicate MSCI [106]. If MSCI does not occur within teleost fish, the X and Y chromosomes of the threespine stickleback would not need to fully synapse outside of the pseudoautosomal region and MSCI would not be triggered. Future work in threespine stickleback should characterize the role of RNA polymerase and repressive histone modifications, like H3K9me3, in regulating the sex chromosomes during meiosis.

Chromosome-wide silencing creates strong constraints on sex-linked genes and inhibits selfish genetic elements from being expressed in meiosis [93, 95, 107, 108]. Sex chromosomes are suspected to be an evolutionary hotspot for the accumulation of selfish genetic elements that can bias chromosomal transmission [95, 107, 108]. We found that X-linked hemizygous genes expressed across all stages of spermatogenesis exhibited higher rates of protein coding evolution. Our findings suggest that X-linked genes involved in spermatogenesis evolve faster than other X-linked genes, evident by higher rates of coding evolution, and expression evolution related to testis specific dosage compensation. Efficient selection of X-linked genes could lead to intragenomic conflicts driven by selfishness between X- and Y- bearing sperm in this species.

Y chromosomes across many species have been found to accumulate gene duplications of testis specific genes [28, 29, 37, 71, 109, 110]. Sex-linked gene amplification has been hypothesized to be the result of selection pressure to escape the repressive environment of silenced chromosomes. Genes escaping post-meiotic repression in mammals have been putatively linked to transmission distortion [26–29, 73, 111, 112]. In the absence of MSCI in the threespine stickleback, we observed that Y-linked duplicates were not enriched in post-meiotic stages. Instead, amplified gene families were expressed throughout meiosis. Y-linked gene duplications on the neo-Y chromosome of *Drosophila miranda* were also found to be expressed during meiosis, rather than being restricted to post-meiotic cell types [82]. Therefore, it may be possible that the enrichment of large ampliconic families explicitly related to post-meiotic expression, may be restricted to mammalian lineages where repression of gene expression extends beyond the meiotic stages of MSCI.

To date, no specific genes have been found to cause segregation distortion in threespine stickleback fish, although two microsatellites have been found to have distorted transmission in an F2 cross [113]. While most lab crosses yield 50:50 sex ratios, some wild populations have been found to be female biased [114]. This could suggest that selection to repress conflicts to maintain even sex ratios is ongoing. Similar to duplicated genes across mammals [29, 111] and *Drosophila* [71, 109, 115], we did find that some duplicated gene families on the threespine stickleback Y chromosome were also expressed in post-meiotic stages. Despite the lack of repression in threespine stickleback, we identified several enriched categories of genes similar to other species, including functions in chromatin assembly and maintenance. The well characterized *Segregation Distorter* meiotic drive system in *Drosophila melanogaster*, distorts transmission by manipulating chromatin state of other cells, leading to sperm dysfunction and near-complete transmission of the *Sd* locus [116]. We also found an enrichment of genes involved in DNA double strand break repair, including elements in the synaptonemal complex, *Smc5.* Y-linked copies of this gene were expressed across both meiotic and post-meiotic stages. Similarly, some of the ampliconic sex-linked gene families in *Mus musculus* (*Slx/Sly*) are orthologs of another synaptonemal complex protein, SYCP3 [117]. Deletions and duplications of *Slx* have been found to alter sex ratios in mice [26]. While the exact mechanisms of how these genes affect transmission are unknown, the role of recombination proteins being duplicated and neo-functionalized across independently derived Y chromosomes is intriguing. Functional testing of the stickleback sex-linked genes may identify a similar role in transmission distortion.

Comparative work across multiple species has showed that several signatures of spermatogenesis have evolved repeatedly, including the amplification of Y-linked spermatogenesis genes, and meiotic/post-meiotic repression. While some features have evolved convergently, many are not conserved across all species [3, 21, 31, 32] The vastly different reproductive strategies and ecological factors affecting fish may alter some of the assumptions made by classic conflict models proposed in mice and flies, and the absence of sex-linked silencing in stickleback fish could be a result of having less selfishness in the male germline.

Most segregation distorter genes have been identified in species with internal fertilization like mice and flies [116, 118, 119]. Externally fertilized eggs in fish could allow for increased male-male competition [120], which may limit the pervasiveness of genomic conflict, due to the deleterious effects on sperm production associated with selfishness [121]. However, reproductive strategies can vary greatly across different species of fish. For example, male stickleback fish court females by building nests, and then provide early stage paternal care by guarding clutches of eggs laid by the female [122], though this behavior does not completely defend against other males [123]. Broader comparisons across teleost fish would provide an interesting approach to study these differences in a comparative framework, as there is variation in the types of mating behaviors, reproductive strategies, and sex chromosomes, across closely related species.

Specifically, it would be noteworthy to compare the pervasiveness of male-male competition to the rate of selfish evolution in spermatogenesis.

## Methods

### Dissociation of testes for single-cell RNA sequencing

Procedures using threespine stickleback fish were approved by the University of Georgia Animal Care and Use Committee (protocol A2024 08-009-Y1-A0).

We performed three runs of single-cell RNA sequencing. Two runs were conducted with a single male fish. The third run was a pool of males (see below). Each male was collected from separate clutches of lab strains derived from wild-caught fish from Port Gardner Bay, Washington. Individuals were selected based on age (∼7 months), testis size, and testis coloration, all of which are indicators that male fish are becoming reproductive and spermatogenesis is proceeding. Stickleback fish have synchronous spermatogenesis. We therefore aimed to select one male that was enriched for cells undergoing meiosis and one male that was enriched for post-meiotic cells. The first male had slightly smaller and lighter colored testes, consistent with earlier meiotic stages, while the second had larger, darker testes consistent with later stages of testis maturation. We sequenced a third pool of two males as a replicate of the first two sequencing runs. We identified two males with different testis coloration and size, which were at different stages of spermatogenesis. Both males were seven-month-old fish collected from separate clutches of fish derived from the same Port Gardner Bay population. We pooled one testis from each male together for tissue dissociation and sequencing.

Cell dissociation was performed separately for each individual. Both testes from each male were placed in ice cold PBS + 0.04% BSA. We followed a previously published single-cell dissociation method used on mouse testes [47]. We first minced the tissue into small pieces with McPherson-Vannas scissors in 200 μL of digestion medium (1mg/mL DNASE, 1mg/mL collagenase/dispace, 1mg/ML hyaluronidase in solution of DMEM/F12). The cell solution was incubated at 37° C for 10 minutes. The cells were filtered through a 40 μm cell strainer to obtain a single-cell suspension, followed by centrifugation at 4° C, 400g for 10 minutes. The pelleted cells were resuspended in 1mL of cold PBS + 0.04% BSA. Cell viability was assessed using trypan blue. From the staining, we determined approximately 90% of the cells were viable. The cells were immediately processed for sequencing using the 10x genomics single-cell sequencing platform with 3’ capture. 10X chromium and subsequent Illumina sequencing were both performed at the Georgia Genome and Bioinformatics Core (University of Georgia).

### Processing of single-cell RNA-sequencing data

The sequencing reads were demultiplexed with CellRanger (v 4.0.0) mkfastq with bcl2fastq (Illumina) and default parameters. We ran CellRanger mkref on the threespine stickleback genome assembly (v5.0) [37, 85]. We aligned fastq reads from all three samples separately using CellRanger count with default parameters, plus the parameter -- chemistry=SC3Pv3. To filter for high quality cells, we used Seurat (v 4.0) to remove cells with > 3000 and <200 unique features (i.e. expressed transcripts) and cells with > 5% mitochondrial counts. Cells with too many or too few identified features likely represent doublets or dead cells, respectively. Transcript expression was normalized using Seurat (v4.0) scaledata, with default parameters. After filtering each dataset separately, we merged all three sequencing runs using Seurat merge, with default parameters. We then performed an analysis on the total dataset, first by identifying cells from each dataset that are contained within nearest neighbors (anchors), by running FindIntegrationAnchors with the dimensions set to dims = 1:20. Then we used Seurat IntegrateData. This step defines a distance between datasets as the total number of cells in the smaller dataset divided by the total number of anchors between each dataset, computes all pairwise distances between each dataset, and clusters the distance matrix to determine a guide tree. After the cells from each dataset were integrated, expression was scaled across each gene, using scaledata. Finally, cells were reassigned to neighbors, clusters, and UMAP was run on the complete dataset.

### Clustering and UMAP

We followed the Seurat (v4.0) workflow [124] to perform cell clustering. We determined dimensionality of the dataset by performing a jackstraw permutation in Seurat. Based on the elbow plot and jackstraw plot (Supplemental Figure 13), we identified 20 significant principal components for the integrated dataset with a P < 0.001. We ran Seurat findneighbors (Dims:1:12) and Seurat findclusters (Resolution=0.8). Graph-based clustering is performed based on construction of a nearest neighbor graph, and then applies a Leiden community detection algorithm. The resolution factor is recommended to be between 0.4-1.2 (https://satijalab.org/seurat/articles/pbmc3k_tutorial.html). We experimented with a range of values from 0.4-1.2 and found that the number of clusters from 0.8 was the most consistent with marker identification and the total number of expected cell types in the threespine stickleback testis [125, 126] To visualize the clusters, we ran the Seurat UMAP function set to 20 dimensions (Dims=1:20). We verified that most clusters had a similar number of reads per cell, and expressed features per cell, to avoid identifying clusters based on read depth alone (Supplemental Figure 14). We also performed alternative clustering approaches by rerunning Seurat findcluster with the added parameter algorithm = Leiden. We then ran scanpy [57] on the counts matrix using default parameters and a clustering resolution of 0.5.

### Identification of cell clusters

Differentially expressed genes were identified using the FindAllMarkers function in Seurat. A Mann-Whitney U test was used to compare gene expression of each gene across all clusters to identify significantly different gene expression profiles between clusters. Genes were considered differentially expressed between clusters if there was a p-value < 0.001 after correcting for multiple comparisons based on a Bonferroni correction. To relate cell clusters to known cell types, we identified upregulated genes specific to clusters that were well characterized genes from mammals [13, 42, 47–50, 127] and fish [51–55].

### Gene ontology

We used the ShinyGO v0.741 web-based tool to identify gene ontology of upregulated genes for each cluster with the genome set to “Stickleback” and processes set to “All available gene sets.” We used all stickleback genes expressed in our single-cell RNA dataset as the background, to identify enrichment of biological processes involved in the significantly up- regulated genes of each cluster. For each set of genes, we calculated the fold enrichment of genes within a pathway and a false discovery rate (FDR). We only retained GO terms with an FDR < 0.01.

### Gene expression across the sex chromosomes

To analyze chromosome-wide expression patterns across cells within a cluster, we extracted alignments from each cluster output using pysam (v 0.16.0.1) and a custom python script (https://github.com/daniel-shaw1/Single-Cell-RNA-sequencing/blob/main/Split_bam_by_cluster.py), that separates RNA alignments for each cluster. The separation of alignment files was run separately for each of the three independent samples based on CellRanger clustering. We counted the total number of reads that aligned within genes for each cluster using htseq-count (v 0.13.5) (-mode union --nonunique all -- minaqual 1 –stranded=no) [128]. These parameters were previously used to identify expression of Y-linked genes in threespine stickleback with bulk RNA-seq [37]. We counted reads within genes of the Y chromosome from published annotations [37] as well as within genes on the autosomes and X chromosome, using annotations from Ensembl (build 107) [129] (Y genes in Supplemental File 2). Reads were counted across the entire transcript as one feature. For both sex chromosomes, we retained all genes with at least a count of one. To assess how gene expression changes on the X and Y chromosomes relative to autosomes across spermatogenesis, we normalized the total expression of each gene on the X and Y by the median expression of all genes across the autosomes of each cluster and then cluster counts were averaged into four categories for each major stage (i.e. somatic, pre-meiotic, meiotic, or post-meiotic).

We also assayed gene expression of duplicate gene families across the Y chromosome that were previously annotated [37]. To analyze the overall expression level of each duplicate gene family, we counted the total number of reads that aligned to each paralog within the family and then summed these counts across all paralogs to get an overall count for each duplicate gene family. We normalized counts for each duplicate gene family per cluster by dividing by the library size of each cluster. Normalized gene duplicate expression was plotted using Morpheus (https://software.broadinstitute.org/morpheus.)

We used previously published bulk RNA-seq from brain [39] (NCBI BioProject PRJNA277770), liver [37] (NCBI BioProject PRJNA591630), and testis [37] (NCBI BioProject PRJNA591630) to identify testis-specific genes expressed from the sex chromosomes in order to distinguish them from genes that were broadly expressed across multiple organs. We aligned three replicates of RNA-seq reads as previously described [66], and identified all of the genes that were only expressed in testis. Genes were considered expressed if they had a read count greater than one.

For identifying allele specific genes in Supplemental Figure 9, we used the full set of sex-linked genes expressed in the testis to confirm allele specific expression of X- and Y- linked genes. We identified all genes that had transcripts aligned to only one sex chromosome in a major cell group in the scRNA-seq, then, we verified mono-allelic expression of those genes in bulk RNA-seq testis to identify a conservative list of mono-allelically expressed genes.

### Detection of retrotransposition throughout the genome

We searched for evidence of retrotransposition of genes from the autosomes and the X chromosome. We aligned all protein coding sequences from the threespine stickleback (NCBI release 100) to the threespine stickleback genome assembly (v. 5) [85]. In order to identify putative retrotranspositions unique to the threespine stickleback lineage after the X and Y chromosomes evolved, we also aligned the threespine stickleback protein coding sequences to an outgroup species, the ninespine stickleback (v. 2.1 genome) [86]. We aligned all protein sequences using Exonerate (v. 2.4) with parameters --model protein2genome --cores 8 --bestn 5 --showalignment FALSE --showvulgar TRUE --ryo “%ps\\n” [130]. We searched for protein sequences that had greater than or equal to one intron that aligned to a second location in the threespine genome with zero introns. These putative retrotranspositions were confirmed to be specific to the threespine stickleback by verifying the alignment did not exist on the orthologous autosome of the ninespine stickleback.

### Protein coding divergence

In order to compare coding sequence evolution, we aligned the ninespine stickleback fish CDS (NCBI NSP_V7) to the threespine stickleback genome [85] using Exonerate (v. 2.4) (exonerate --model est2genome -M 2000 --bestn 15 --percent 25) [130]. We ran codeml from the PAML (phylogenetic analysis by maximum likelihood) (runmode =-2) software package [131] to calculate *d*_N_, *d*_S_, and *d*_N_/*d*_S_ for all alignable coding sequence between the threespine and ninespine stickleback. Exonerate alignments were filtered to remove stop codons and gaps, and processed into a codeml compatible input using a custom Perl script (https://github.com/daniel-shaw1/Single-Cell-RNA-sequencing), modified from a previous study [37]. For genes with multiple annotated transcripts that aligned between the threespine and ninespine stickleback, *d*_N_/*d*_S_ ratios were averaged across the transcripts. Genes with a *d*_N_/*d*_S_ value of 99 were removed as these values are unreliable and occur when a gene does not have synonymous substitutions. For genes with a *d*_N_/*d*_S_ estimate, we identified genes that had higher expression in one of the major cell types if they had 2x the expression of the median number of counts across the other three groups. These counts were found using a custom R script (https://github.com/daniel-shaw1/Single-Cell-RNA-sequencing/blob/main/induced_dnds.r.)

### Meiotic chromosome spreads

The methods used for chromosome spreads were adapted from a protocol for zebrafish [132]. We collected fish that were undergoing meiosis that were five to eight months in age and had minimal pigmentation of the testes [41]. Whole testes were removed and submerged in 1 mL of 1X PBS inside a 1 mL Douce homogenizer. The tissue was homogenized by grinding with the tight pestle approximately 10 times. The entire volume of cell suspension was transferred to a 1.6 mL tube and centrifuged at 500 xg for five minutes. The supernatant was then removed and the pellet was resuspended in 600 μL of 0.8% sodium citrate. The volume of sodium citrate was adjusted based on the size of the pellet to achieve a low turbidity solution. The sodium citrate cell suspension was incubated at room temperature for five to eight minutes. 20 μL of the cell suspension and 20 μL of 100mM sucrose was pipetted onto each slide and spread by gently tilting the slide. 100 μL of 1% (pH 8.5, adjusted with 0.1 M sodium tetraborate) paraformaldehyde solution with 0.15% TX-100 was then added and spread across each slide before incubating in a humid chamber at room temperature for two and a half hours. After incubating the slides, we removed the top of the humid chamber to cure the slides overnight.

After curing, the slides were washed 3 consecutive times for 15, five, and five minutes in 1X PBS with gentle agitation. Slides were allowed to dry for 15-20 minutes before immunofluorescence or storing at –20°C.

### Immunofluorescence staining of SMC3 and γH2AX

1.5 mL of permeabilization solution (1X PBS, 1 mM EDTA, and 1% TritonX-100) was pipetted onto the slides and incubated for 20 minutes at room temperature. After dumping off the solution, the slides were blocked with 1.5 mL of 1X antibody dilution buffer, incubating at room temperature for 30 minutes. A 1:80 dilution of primary antibodies in 40 μL total volume was pipetted on each slide (anti-SMC3, Abcam ab9263; anti- γH2AX, DSHB UNC93-5.2.1). The slides were sealed with coverslips and rubber cement before being incubated in a humid chamber overnight at 4°C. After incubating overnight, the coverslips were allowed to float off in 1X PBS, and the slides were blocked with 1.5 mL 1X antibody dilution buffer for 30 minutes at room temperature. A 1:100 dilution of secondary antibodies in 40 μL total volume was pipetted on each slide (goat anti-rabbit Alexa Fluor 488, Abcam ab150077; goat anti-mouse Alexa 568, ThermoFisher A-21134). The slides were sealed with coverslips and rubber cement before incubating in a humid chamber at 37°C for two to three hours. After the coverslips were removed, the slides were washed three times for 15, five and five minutes in 1X PBS with gentle agitation. 40 μL of Vectashield Antifade Mounting Media with DAPI (Vector Labs) was added to the slides and sealed with coverslips and nail polish.

### Staging of Meiocytes and Super Resolution Imaging

The chromosome spreads were viewed and staged with a Nikon Eclipse Ti2 Microscope with a 100x oil immersion lens. Prophase I stages were identified with the following criteria: small, non-contiguous stretches of SMC3 were defined as leptotene; long SMC3 pieces and incomplete strands indicating an absence of synapsis were defined as zygotene; and pachytene was identified by fully formed SMC3 strands, suggesting complete synapsis. Final images were taken with a Zeiss Elyra S1 (SR-SIM) Super Resolution Microscope with 100x oil immersion lens. The images were processed with ImageJ and Zeiss Black software.

## Data availability

The single cell data for this manuscript is available within the short reads archive under PRJNA1191497. The Bulk RNA-seq is previously published and available on the short reads archive: Brain: PRJNA277770, Liver: PRJNA591630, Testis: PRJNA591630.

Scripts and bioinformatic pipelines are available at: https://github.com/daniel-shaw1/Single-Cell-RNA-sequencing

All microscopy images are available upon request.

## Acknowledgements

This work was supported by the National Science Foundation MCB 1943283 (M.A.W.), the National Institutes of Health R01GM147312 (M.A.W.), the University of Georgia Research Foundation (D.E.S.), and by the Society for Study of Evolution R.C. Lewontin Early Award (D.E.S). Single cell library preparations and sequencing were performed at the UGA Georgia Genome and Bioinformatics Core.

---

**Supplemental Figure 1.**
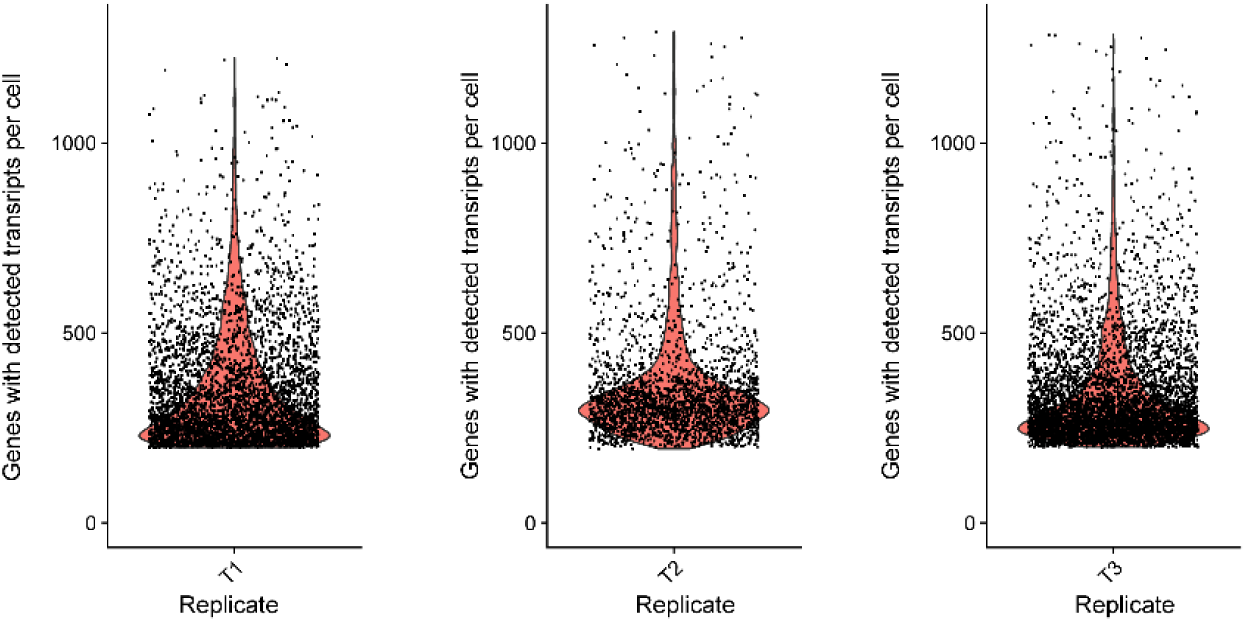
Gene counts per samples. Number of transcripts detected for each gene was totaled for each cell barcode within each of the three single cell RNA sequencing experiments. A similar distribution of detected genes were identified in each of three experiments.

**Supplemental Figure 2.**
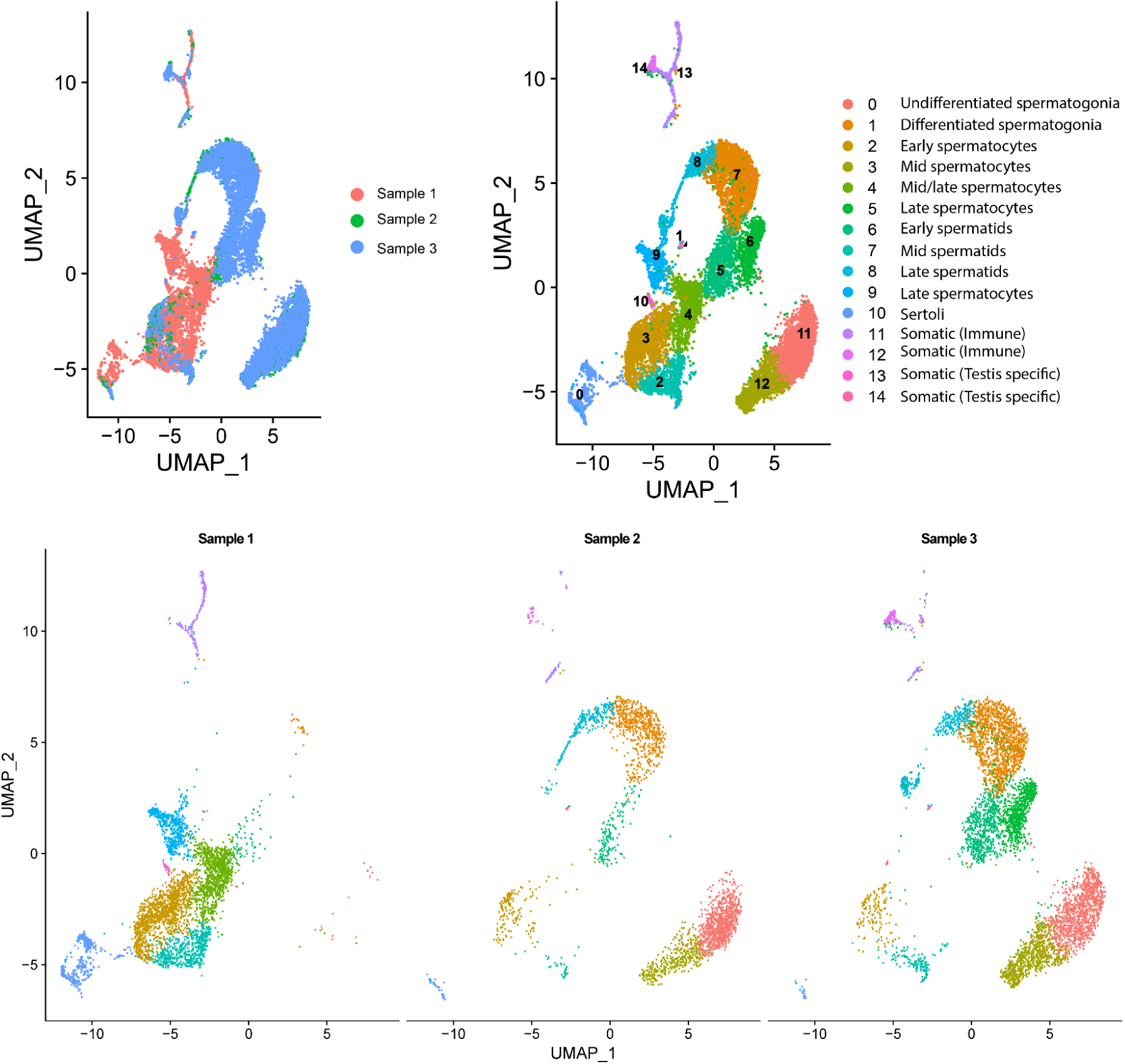
Clusters split by sequencing replicate. Cell barcodes from all three sequencing runs were merged, and UMAP is depicted for all merged cells (top), and for each replicate separately (bottom). Proportion of each cell type within each sample was dependent on the onset of synchronous spermatogenesis in each individual. Only cell clusters identified in two of three replicates were used for downstream analysis.

**Supplemental Figure 3.**
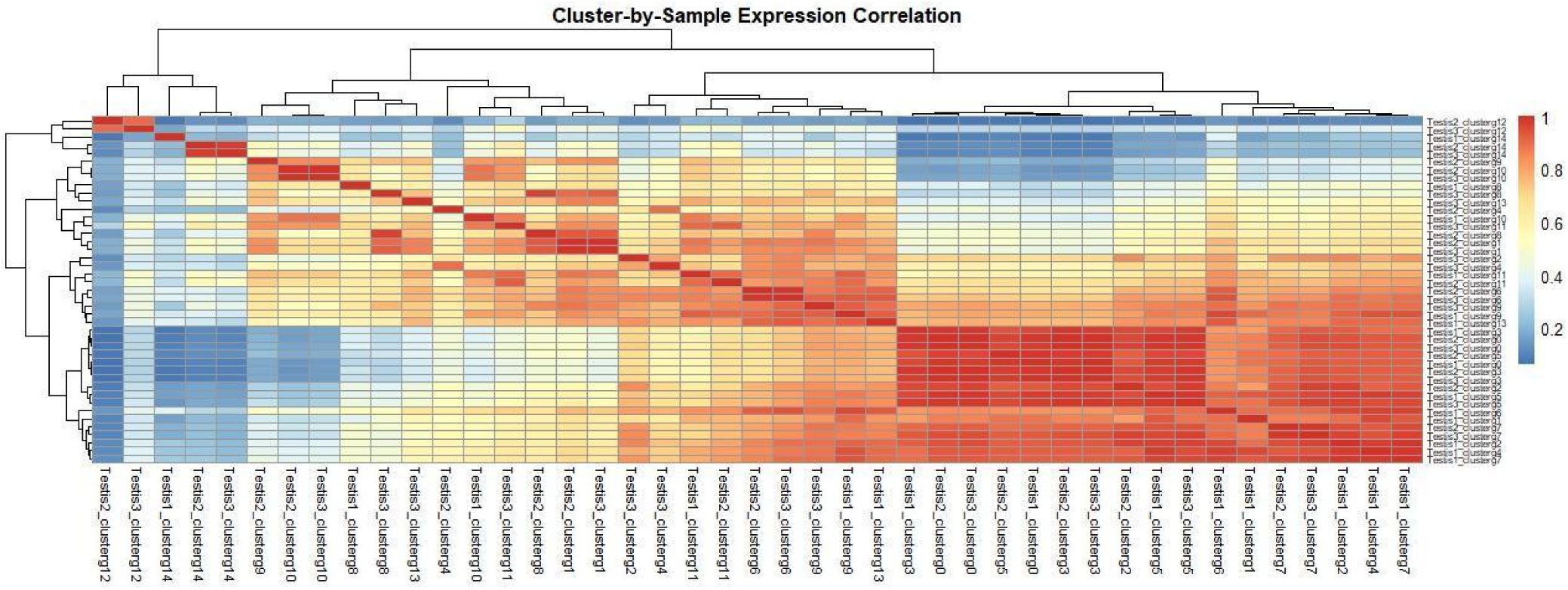
Cluster wide expression correlation across all samples. Normalized gene expression of the top 500 expressed genes in each cluster was averaged and compared for each cluster and sample. Hierarchical clustering reveals that most clusters cluster by cell types across replicates rather than by sample.

**Supplemental Figure 4.**
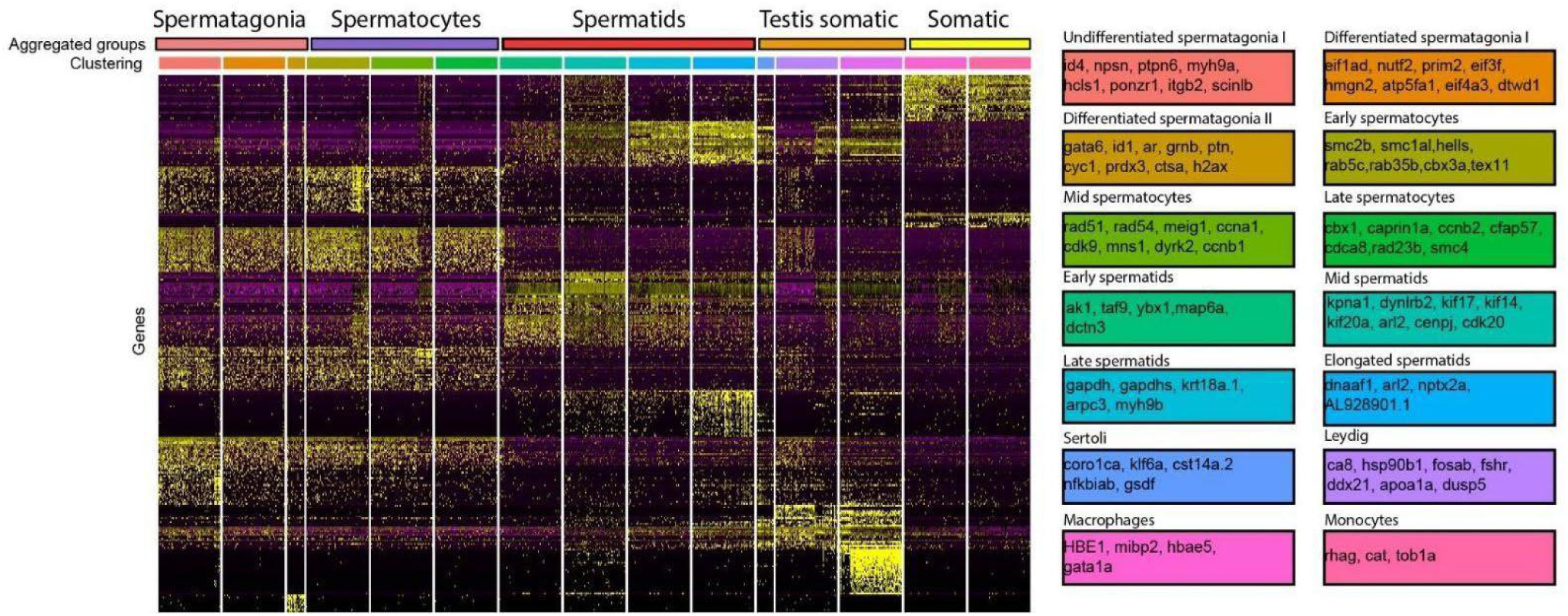
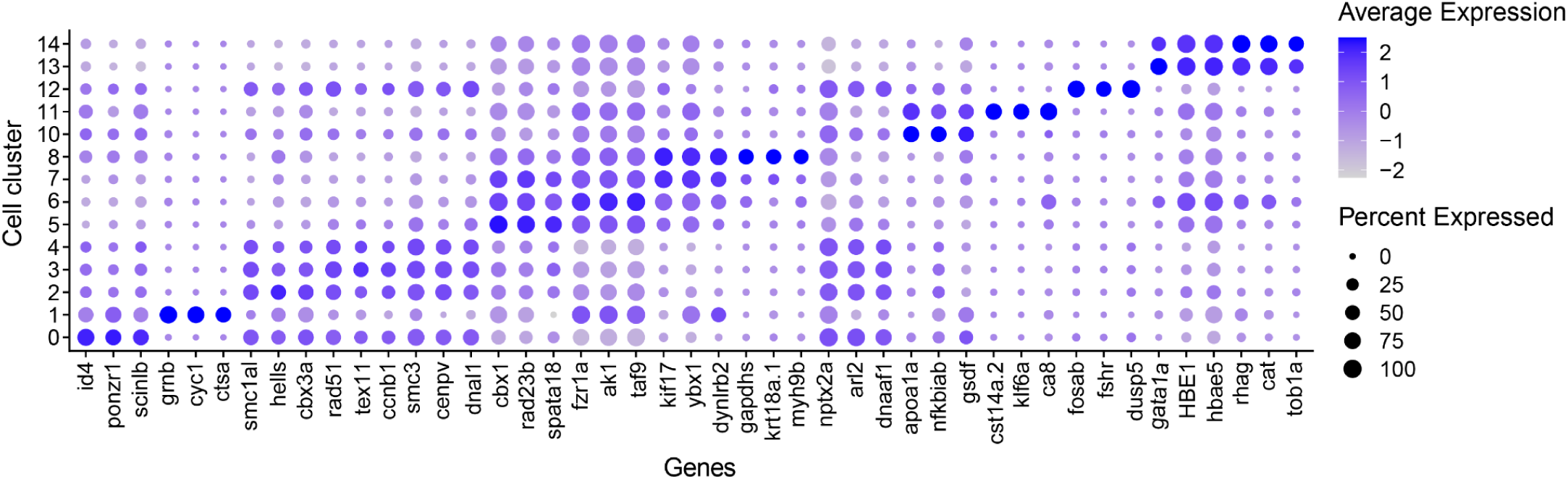
Marker genes for all clusters. Differentially expressed genes in each cell cluster correspond with diverse testis cell types. Heatmap was generated for the top 250 variable expressed genes across all clusters. Represented marker genes for each cluster are listed in boxes. Clusters were assigned to five major cell groups for downstream analysis.based on similar expression and biological functions. Bottom: Cluster dotplots for representative marker genes. Marker genes were identified based on having significantly different expression (P < 0.01, Mann-Whitney U test), and had a confirmed function in the cell type in other species.

**Supplemental Figure 5.**
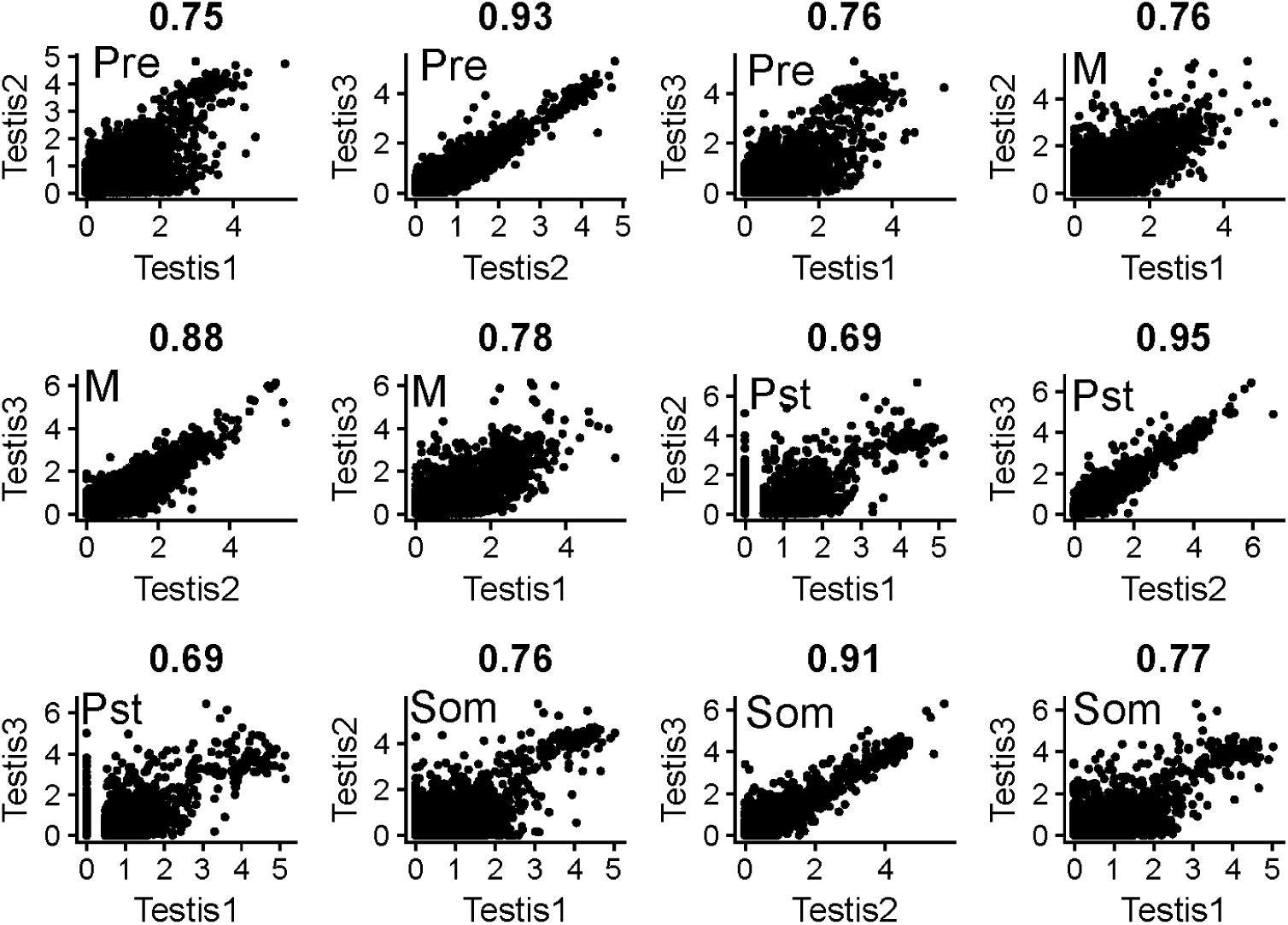
Correlation of cluster expression profiles across replicates. Gene expression for all expressed genes identified in clusters were highly correlated (R > 0.69) across samples. Testis 2 and 3 showed stronger correlations across cell types compared to s, likely due to a similar developmental staging at the time of testis dissociation. Spearman rank correlation is reported above each graph. Cell types are noted as Pre (pre-meiotic), M (meiotic), Pst (post-meiotic), and Som (somatic).

**Supplemental Figure 6.**
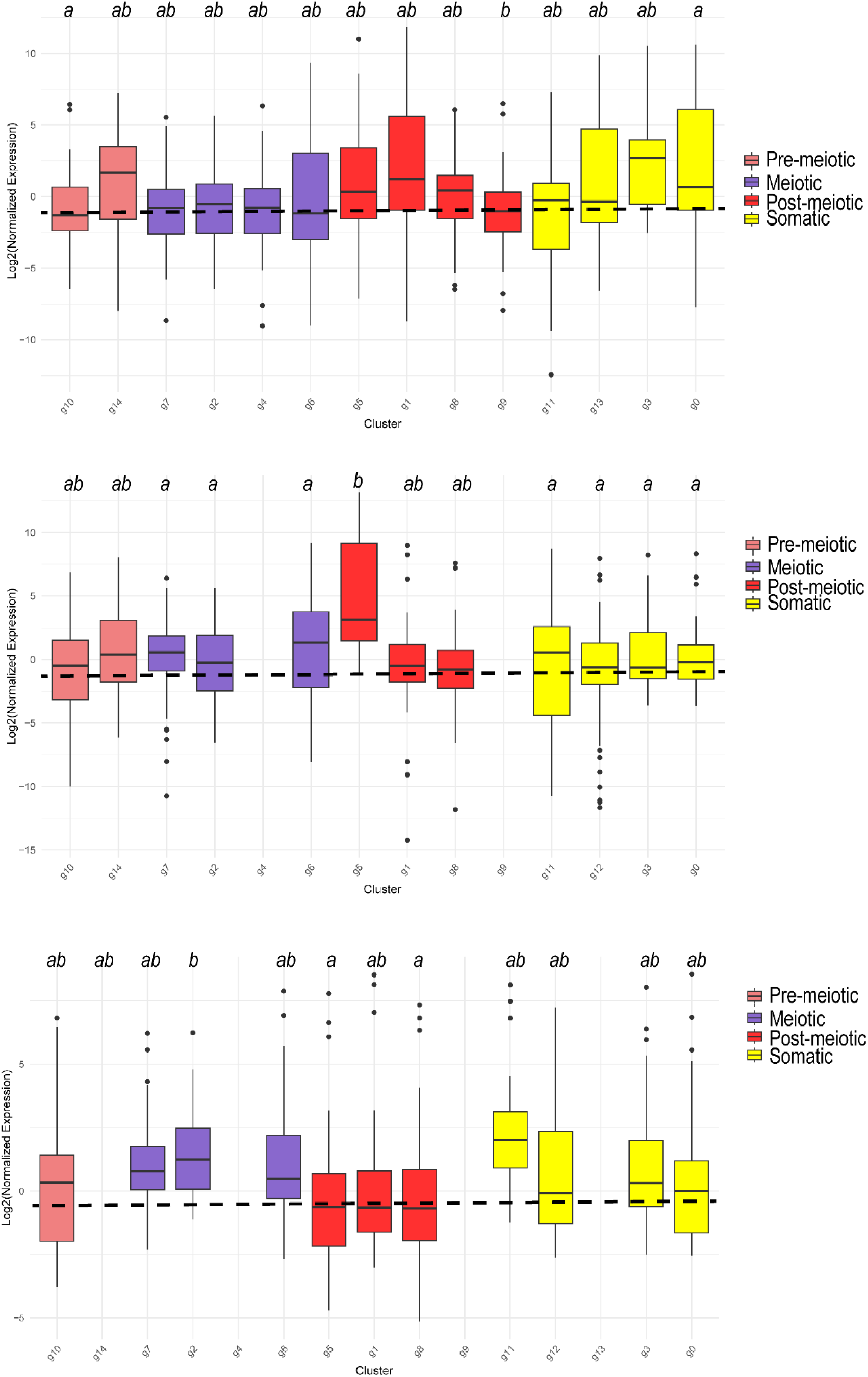
X:AA expression across all clusters and samples. No evidence for MSCI across all three samples when examining all identified clusters. X gene expression was normalized by autosomal gene expression for each cluster across all three samples. Letters denote statistical significance based on a Kruskal Wallis test and Dunn’s test. Dotted line represents median X:A expression level across main cell groups in Figure 2.

**Supplemental Figure 7.**
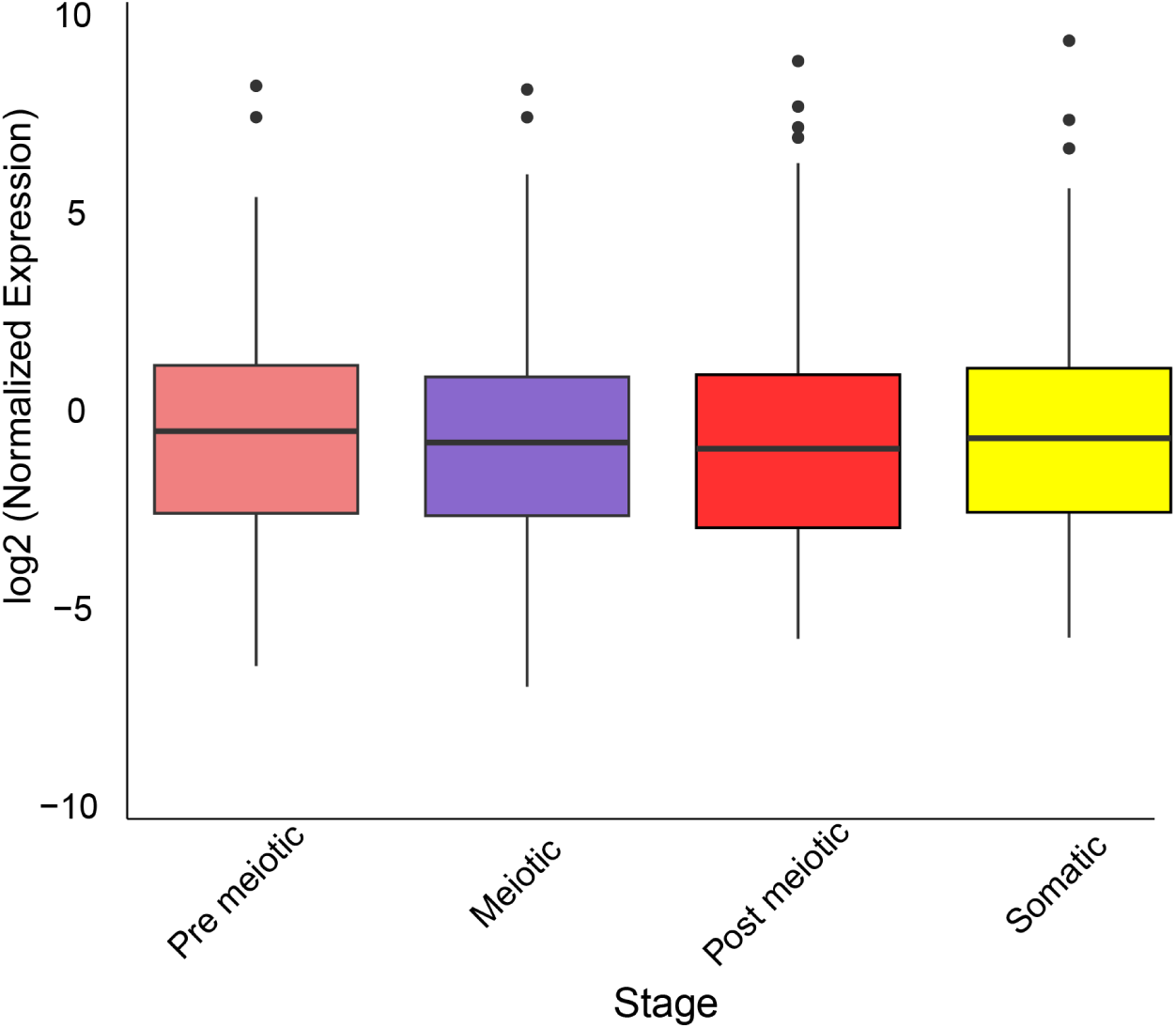
Broadly expressed genes are maintained across all stages of spermatogenesis. Bulk RNA-seq data from three tissues brain, liver, and testis were compared to identify genes that were expressed testis specifically. When we filtered out 20 testis specific X-linked genes, we still identify similar expression across all stages of spermatogenesis P > 0.05 for all comparisons, Kruskal-Wallis and Dunn’s test.

**Supplemental Figure 8.**
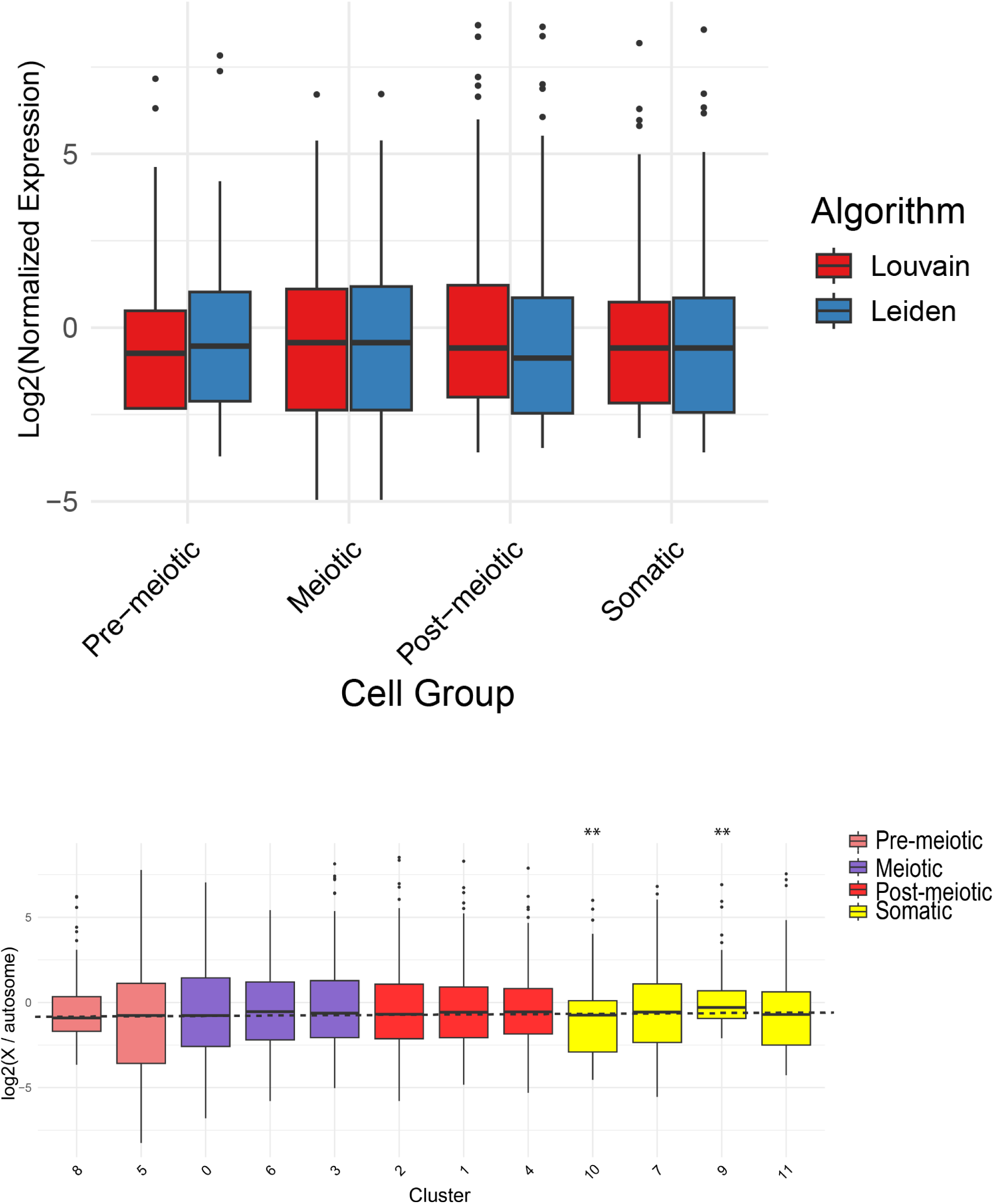
Lack of MSCI is robust across distinct cell clustering algorithms and software. Top: Normalized X expression was compared between two distinct clustering algorithms in Seurat. X expression was similar across major cell groups for both algorithms (P = 0.441, Kruskal Wallis test. Bottom: Cell clustering was reanalyzed with a separate software package (scanpy) and re-annotated using cell markers. Scanpy clustering was performed using all three samples. Normalized X expression is similar across all stages of spermatogenesis. Only two somatic clusters had variable expression (P < 0.05, Kruskal wallis and Dunn’s test).

**Supplemental Figure 9.**
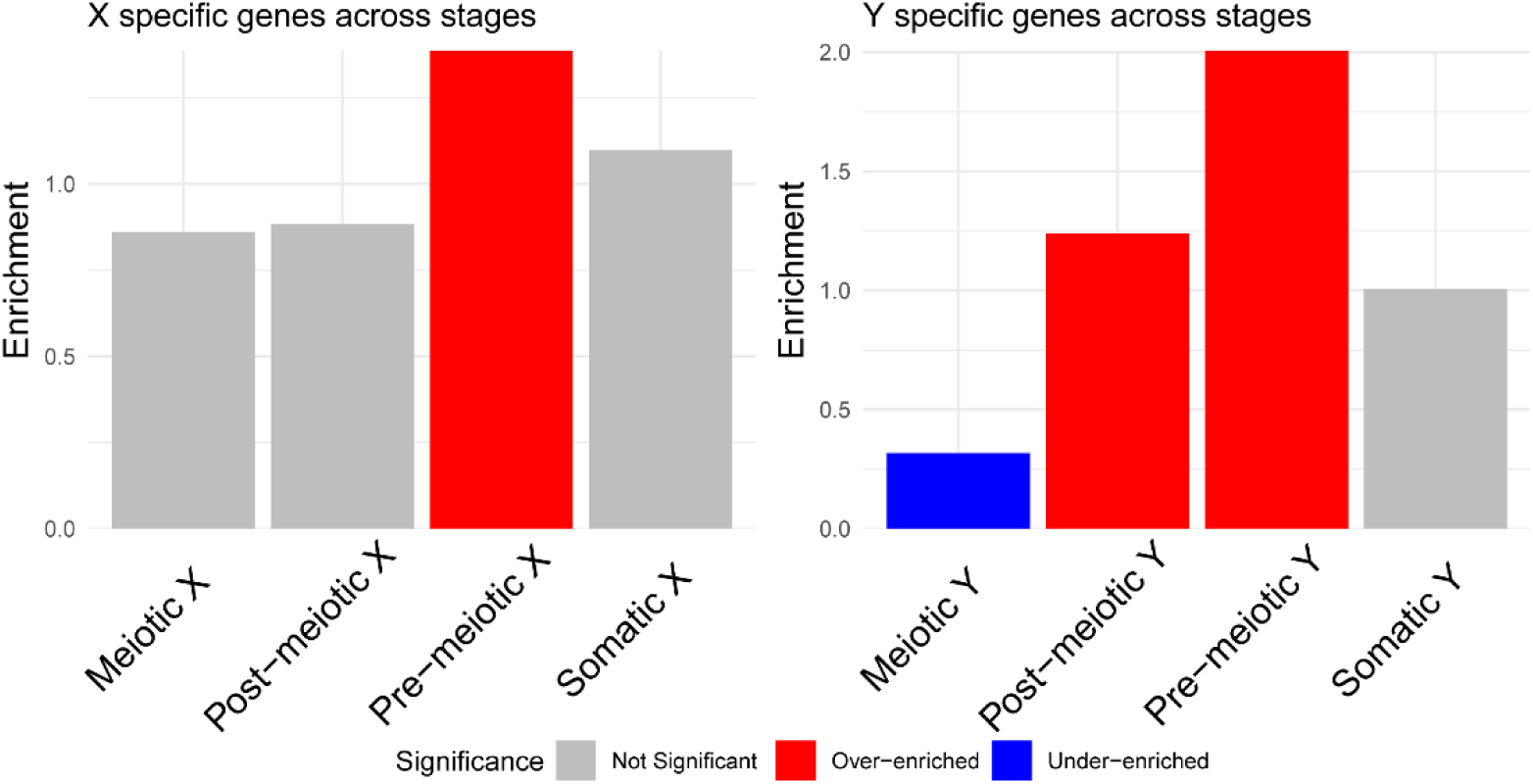
Enrichment of X- and Y- mono-allelic expressed genes across stages of spermatogenesis. Gametolog specific expression was identified for the 4 main cell types of spermatogenesis by aggregated counts from all clusters. Genes with transcripts aligned to only one sex chromosome per cluster were identified. Mono-allelic expression was confirmed using bulk RNA-seq from three replicates of meiotically active testis. Enrichment of mono- allelic expression across cell stages was calculated using a hypergeometirc test comparing the number of mono-allelic expressed genes observed to expected based on the number of genes expressed in that cell type. X- only expression was overenriched in pre-meiotic cells (P < 0.01). Y-only expression was over-enriched in pre-meiotic and post-meiotic cell types (P < 0.01), and under-enriched in meiotic cell types (P < 0.01).

**Supplemental Figure 10.**
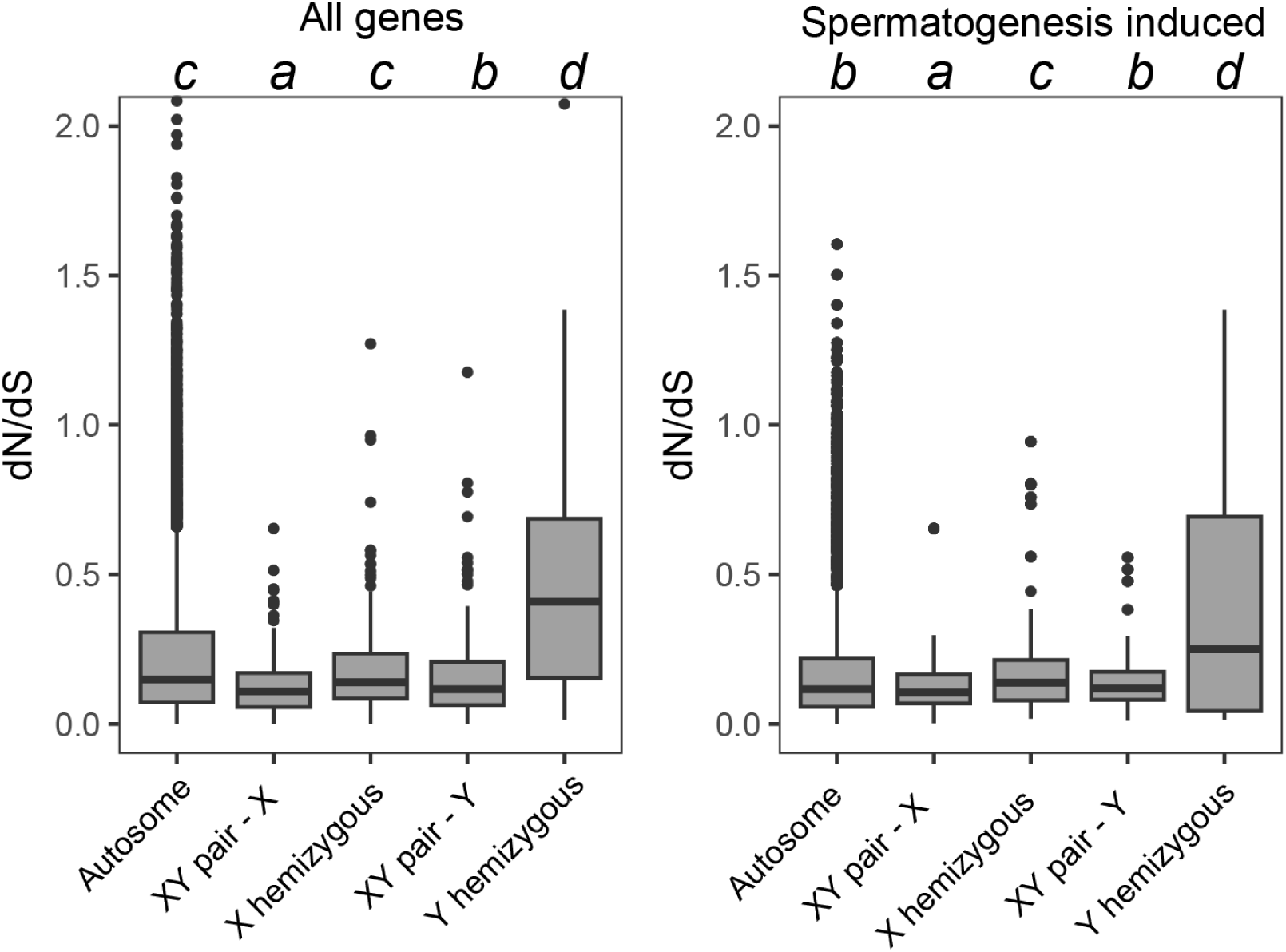
d_N_/*d*_S_ across all induced genes in major cell groups of spermatogenesis. *d*_N_/*d*_S_ was calculated between ninespine and threespine stickleback for genes on autosomes, X hemizygous genes, X and Y gametologs, and Y homozygous genes. scRNA-seq was used to identify genes that were induced in four major cell groups. Letters denote statistical comparisons across all remaining stages and categories based on Kruskal Wallis and Dunn’s test. Autosomes have the highest count with 2,262 genes followed by X hemizygous genes with 50 genes, X gametologs with 31 genes, Y gametologs with 31 genes, and Y hemizygous genes with just 4 genes.

**Supplemental Figure 11.**
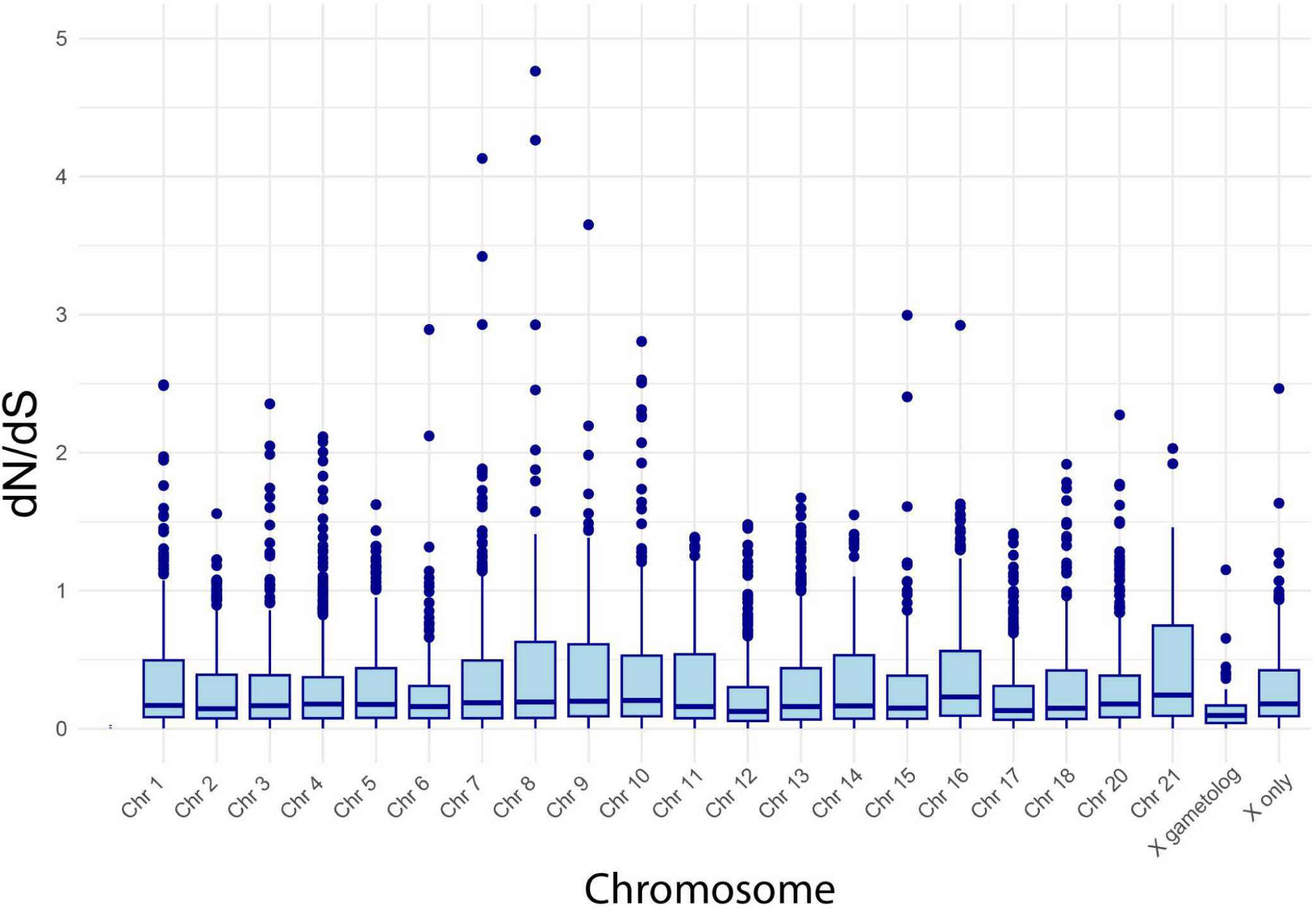
d_N_/*d*_S_ between threespine and ninespine stickleback for all chromosomes. Both X gametologs and Chr 12 (Ninespine sex chromosomes) have lower dN/dS than median autosomal (P < 0.01, Kruskal Wallis and Dunn’s test). All other comparisons are not statistically different (P > 0.05 for all pairwise comparisons).

**Supplemental Figure 12.**
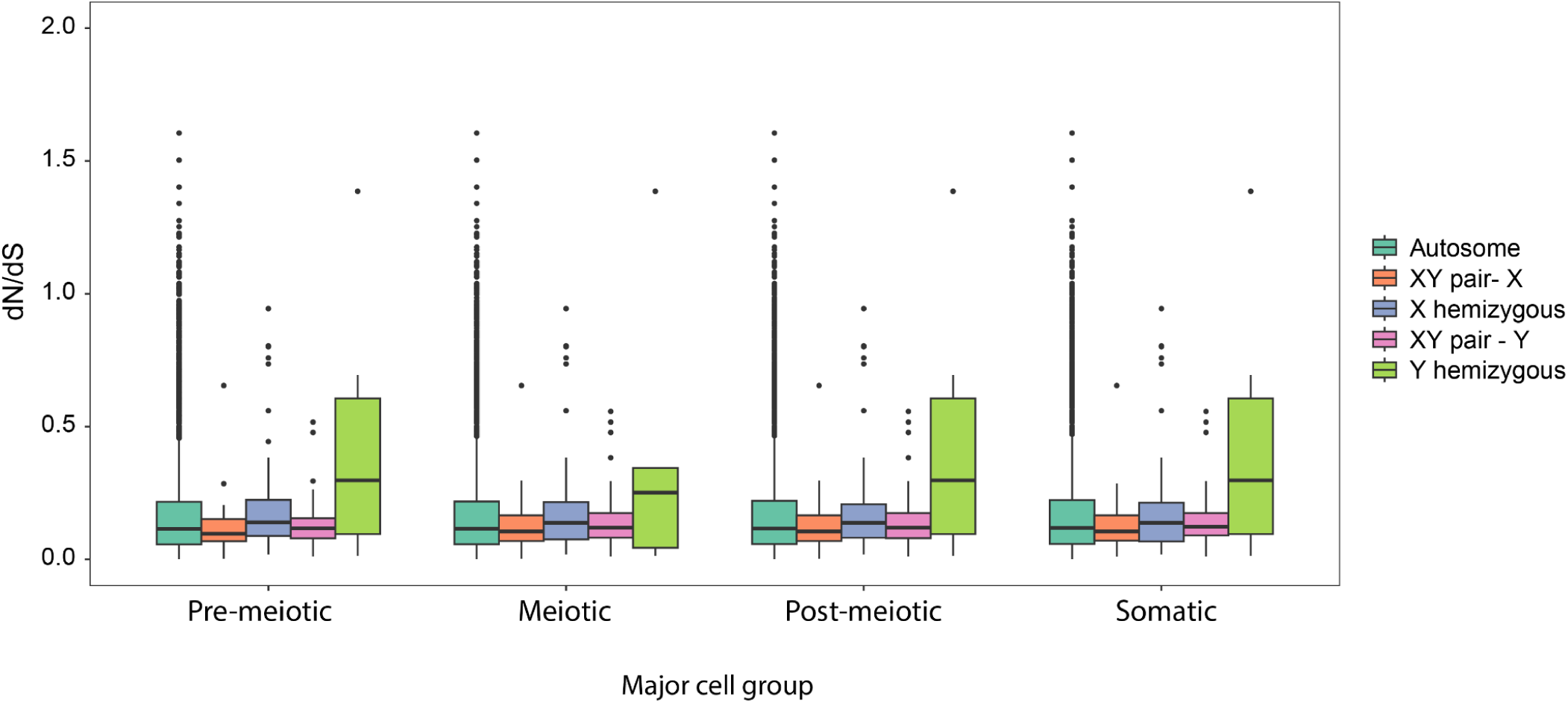
Similar signatures of selection across spermatogenesis. *d*_N_/*d*_S_ was calculated between ninespine and threespine stickleback for genes on autosomes, X hemizygous genes, X and Y gametologs, and Y homozygous genes. scRNA-seq was used to identify 2378 genes that were induced in four major cell groups. There were no significant differences between stages for each chromosome group (P = 0.501 Kruskal Wallis and Dunn’s test). The dataset includes genes that were induced and had an alignment between threespine and ninespine stickleback cross four developmental stages for each chromosome category. Autosomes have the largest sample sizes, with 414, 465, 340, and 1,043 genes across the four stages, respectively. X gametologs include 16, 3, 5, and 7 genes; X-only genes have 24, 9, 9, and 8 genes; Y gametologs consist of 16, 3, 5, and 7 genes; and Y-only genes are the smallest group, with just 2 genes in each of the last two stages.

**Supplemental Figure 13.**
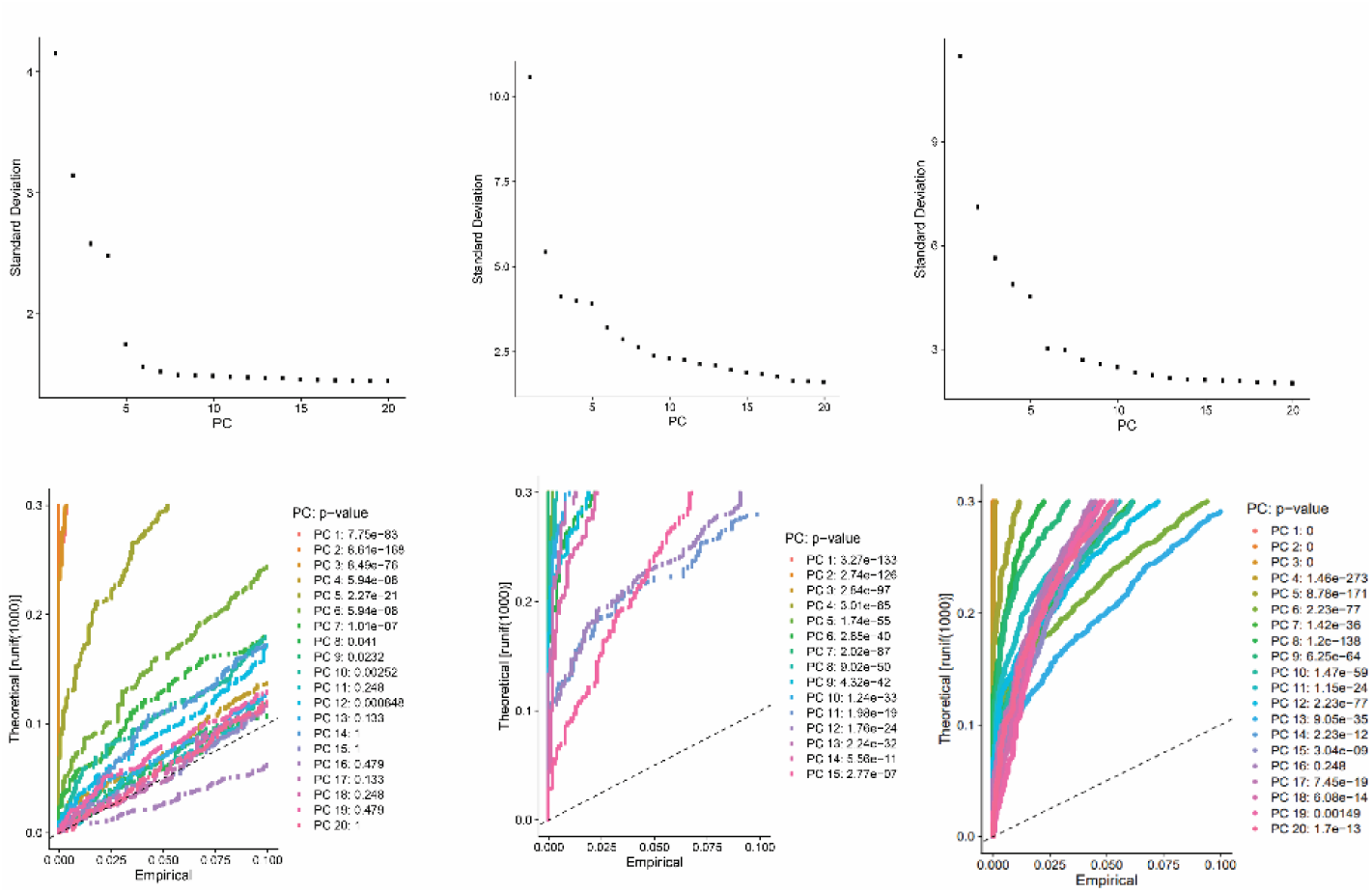
Identification of significant principal components for scRNA-seq. Elbow plots and jackstraw simulations were generated to assess the number of significant principle components for each sequencing experiment.

**Supplemental Figure 14.**
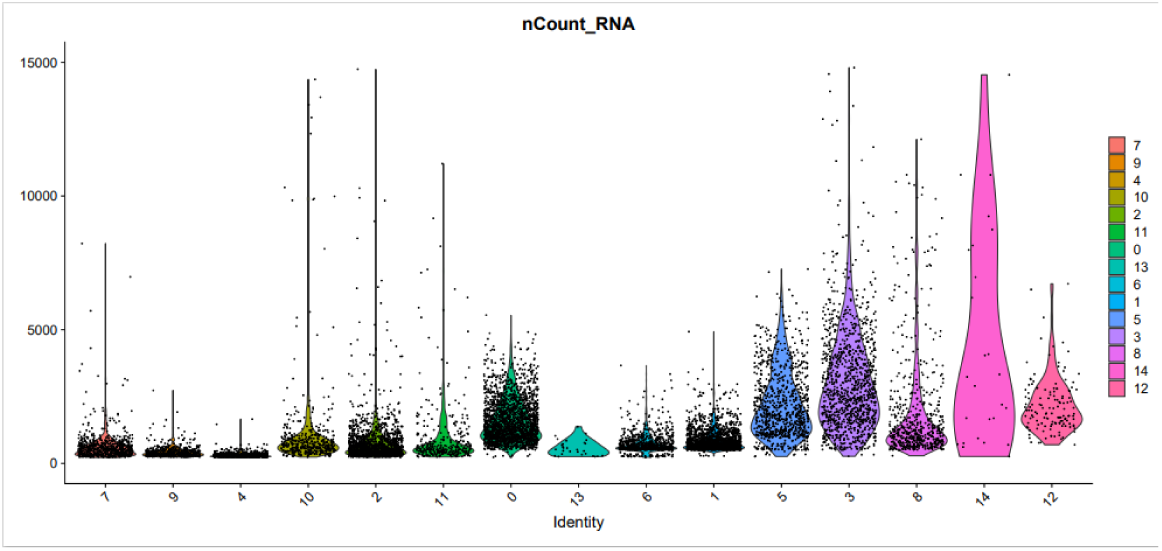
Number of reads per cell type. Number of RNA molecules identified per cell within each cell cluster were counted using Seurat.

**Supplemental File 1. List of all differentially expressed genes identified for each cell type.**

**Supplemental File 2. List of all annotated Y genes and coordinates.**

