## Supplemental Figures for "Single cell RNA-sequencing reveals no evidence for meiotic sex chromosome inactivation in the threespine stickleback fish"

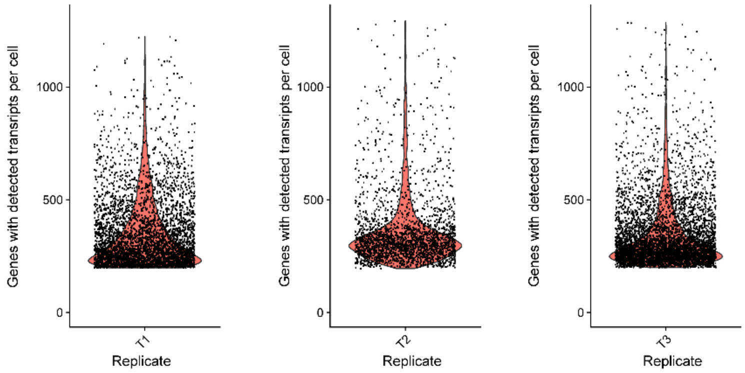


**Supplemental Figure 1. Gene counts per samples.**

Number of transcripts detected for each gene was totaled for each cell barcode within each of the three single cell RNA sequencing experiments. A similar distribution of detected genes were identified in each of three experiments.

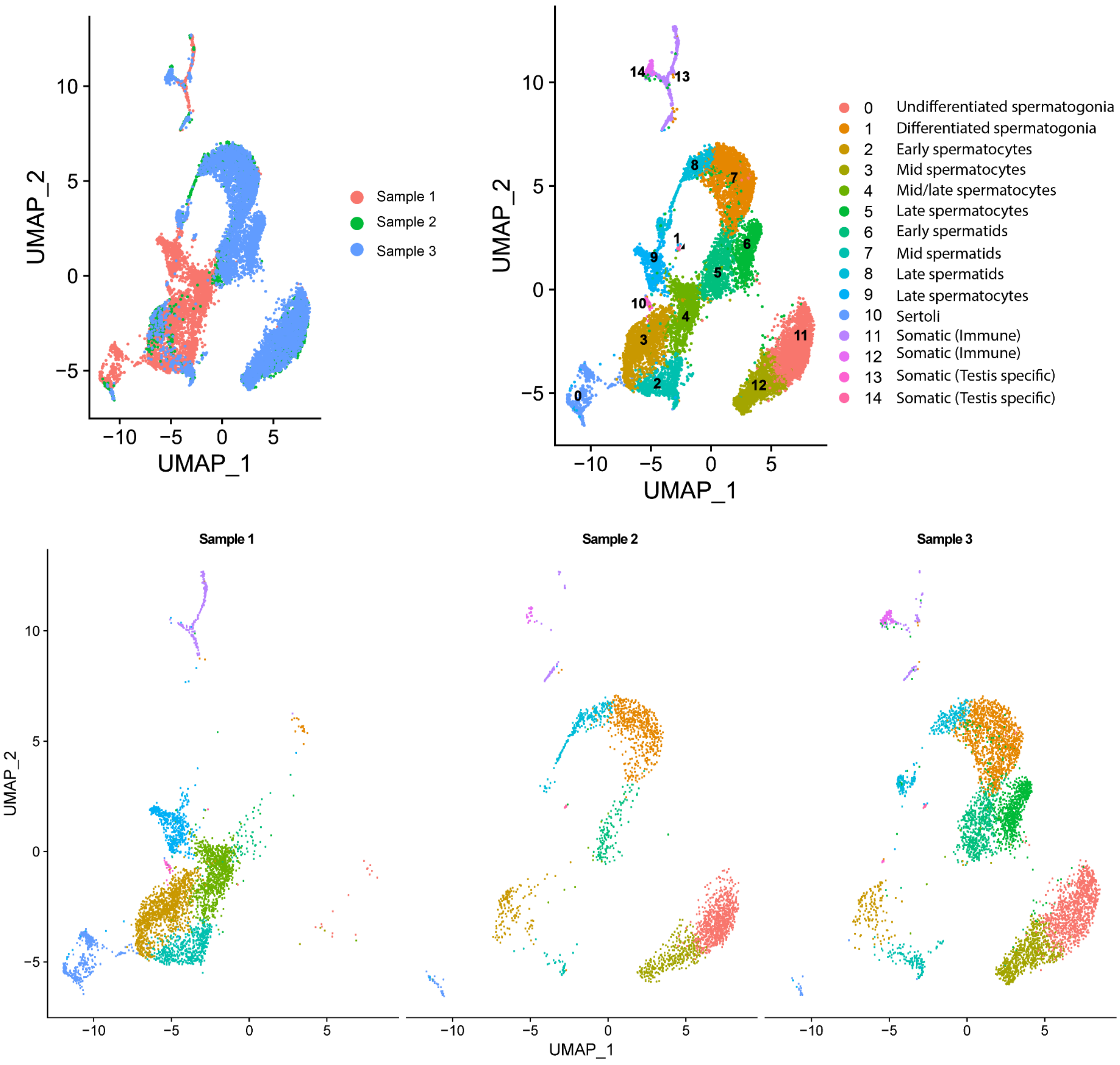


**Supplemental Figure 2. Clusters split by sequencing replicate.** Cell barcodes from all three sequencing runs were merged, and UMAP is depicted for all merged cells (top), and for each replicate separately (bottom). Proportion of each cell type within each sample was dependent on the onset of synchronous spermatogenesis in each individual. Only cell clusters identified in two of three replicates were used for downstream analysis.

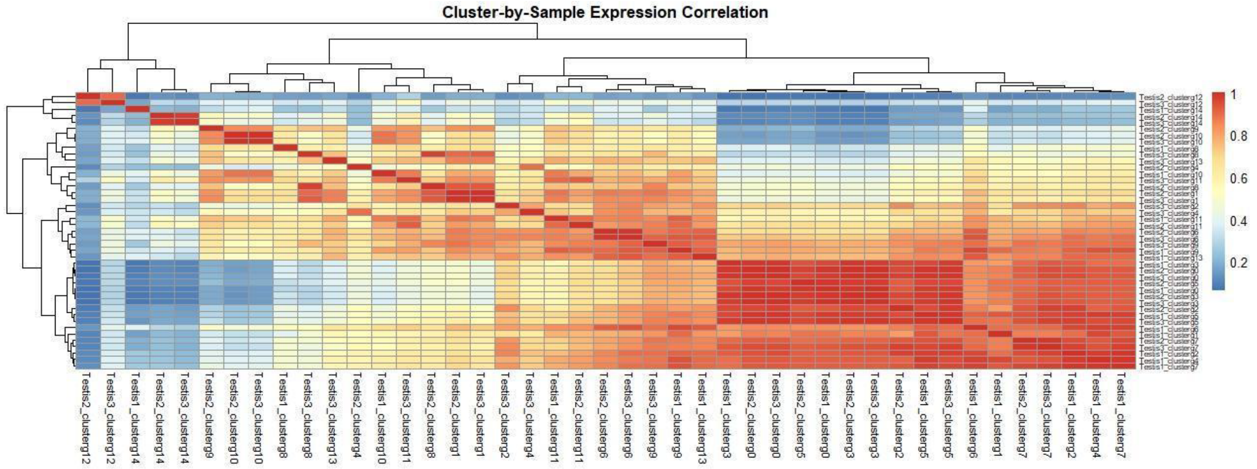


**Supplemental Figure 3. Cluster wide expression correlation across all samples.** Normalized gene expression of the top 500 expressed genes in each cluster was averaged and compared for each cluster and sample. Hierarchical clustering reveals that most clusters cluster by cell types across replicates rather than by sample.

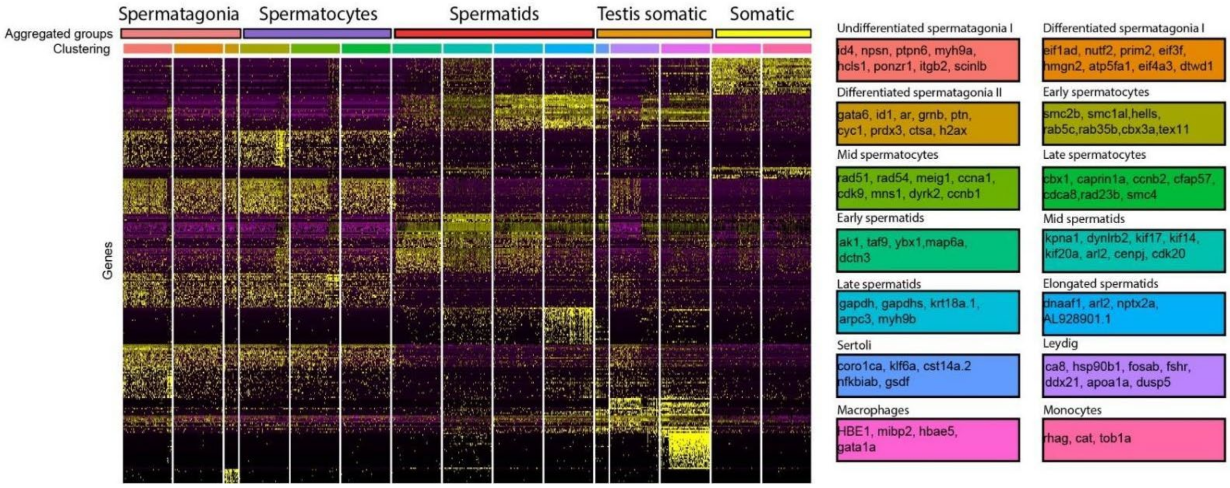

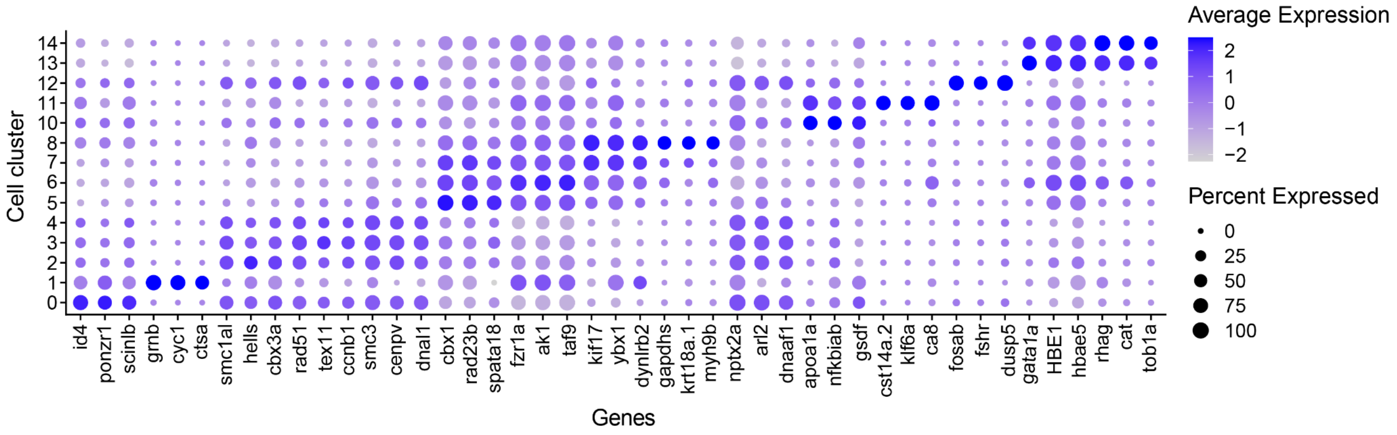


**Supplemental Figure 4. Marker genes for all clusters.** Differentially expressed genes in each cell cluster correspond with diverse testis cell types. Heatmap was generated for the top 250 variable expressed genes across all clusters. Represented marker genes for each cluster are listed in boxes. Clusters were assigned to five major cell groups for downstream analysis.based on similar expression and biological functions . Bottom: Cluster dotplots for representative marker genes. Marker genes were identified based on having significantly different expression ($P < 0.01$, Mann-Whitney U test), and had a confirmed function in the cell type in other species.

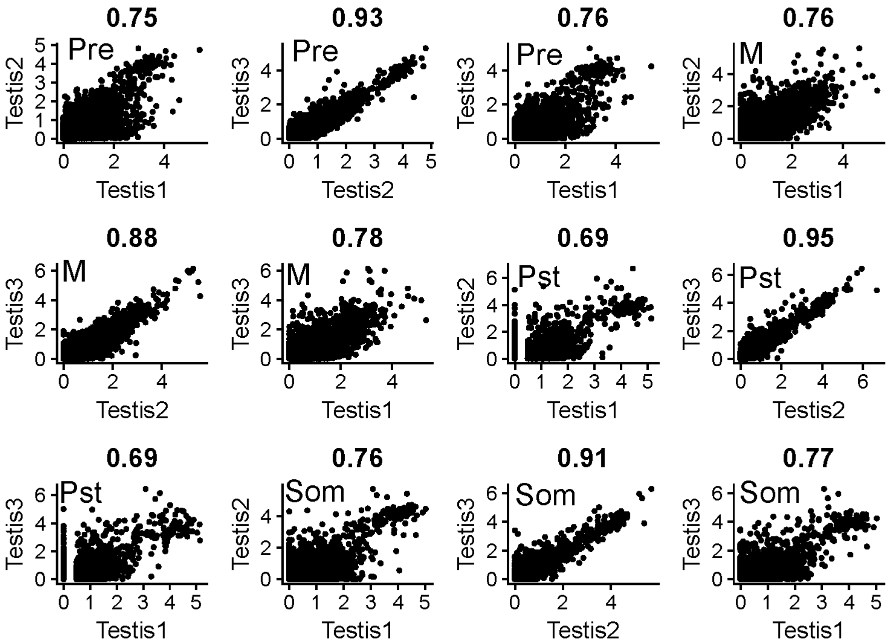


**Supplemental Figure 5. Correlation of cluster expression profiles across replicates**

Gene expression for all expressed genes identified in clusters were highly correlated (R > 0.69) across samples. Testis 2 and 3 showed stronger correlations across cell types compared to s, likely due to a similar developmental staging at the time of testis dissociation. Spearman rank correlation is reported above each graph. Cell types are noted as Pre (pre-meiotic), M (meiotic), Pst (post-meiotic), and Som (somatic).

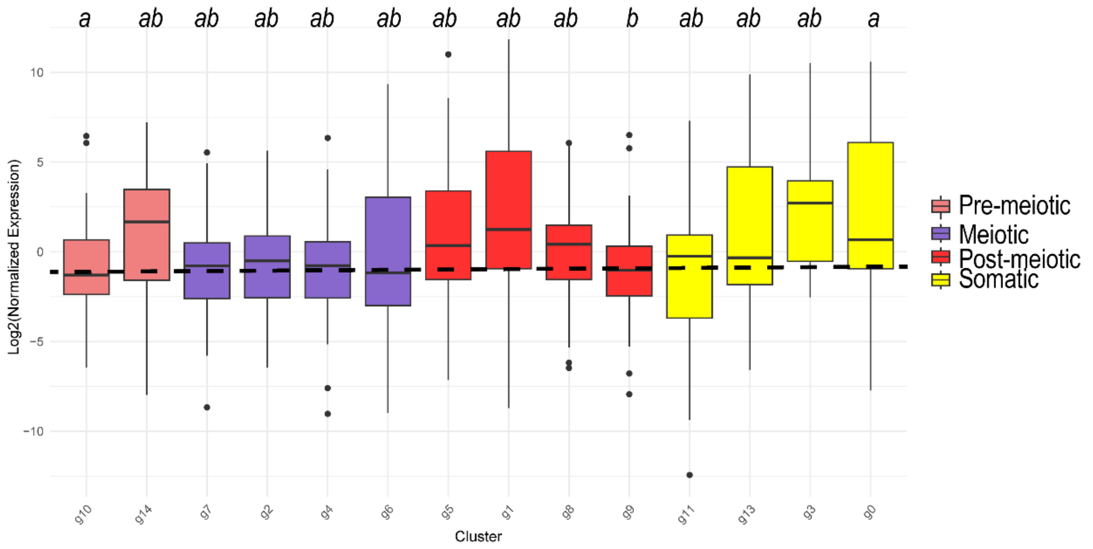
a ab ab ab ab ab ab ab ab b ab ab ab a
Log2(Normalized Expression)
10
5
0
−5
−10
g10 g14 g7 g2 g4 g6 g5 g1 g8 g9 g11 g13 g3 g0
Cluster
Pre-meiotic
Meiotic
Post-meiotic
Somatic

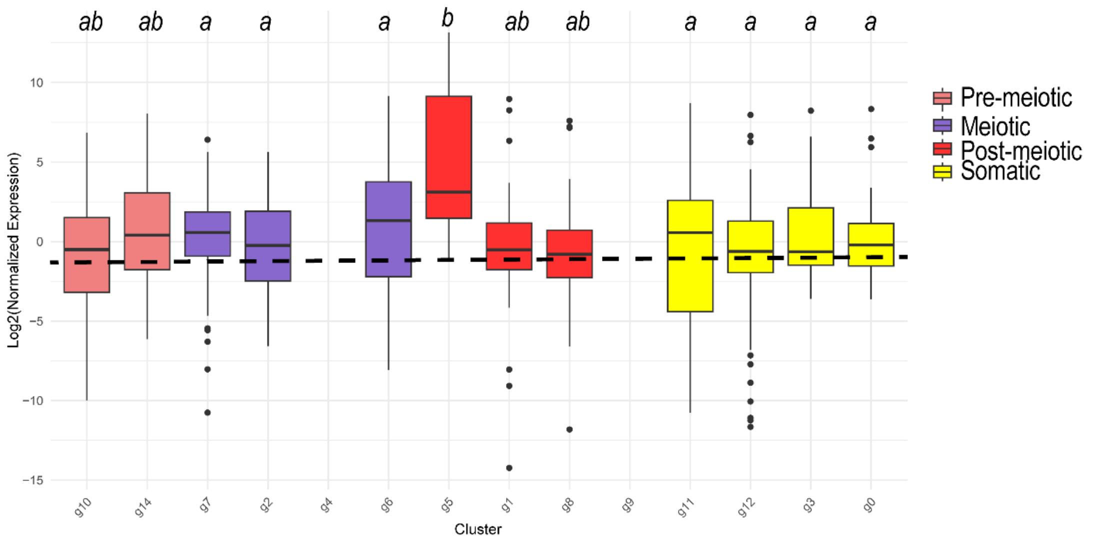
ab ab a a a b ab ab a a a a
Log2(Normalized Expression)
10
5
0
−5
−10
−15
g10 g14 g7 g2 g4 g6 g5 g1 g8 g9 g11 g12 g3 g0
Cluster
Pre-meiotic
Meiotic
Post-meiotic
Somatic

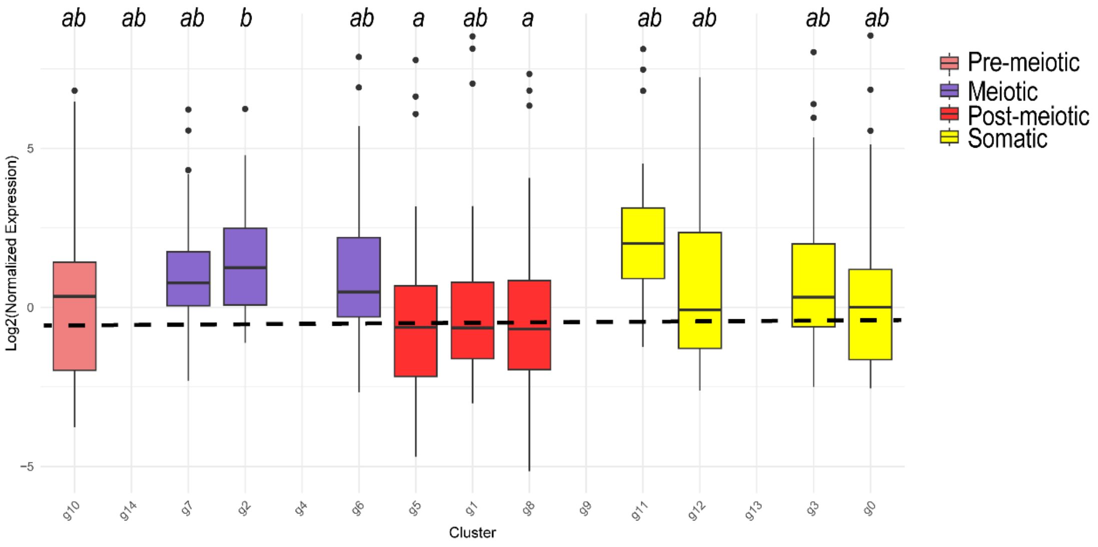
ab ab ab b ab a ab a ab ab ab ab
Log2(Normalized Expression)
5
0
−5
g10 g14 g7 g2 g4 g6 g5 g1 g8 g9 g11 g12 g13 g3 g0
Cluster
Pre-meiotic
Meiotic
Post-meiotic
Somatic

**Supplemental Figure 6. X:AA expression across all clusters and samples**

No evidence for MSCI across all three samples when examining all identified clusters. X gene expression was normalized by autosomal gene expression for each cluster across all three samples. Letters denote statistical significance based on a Kruskal Wallis test and Dunn's test. Dotted line represents median X:A expression level across main cell groups in Figure 2.

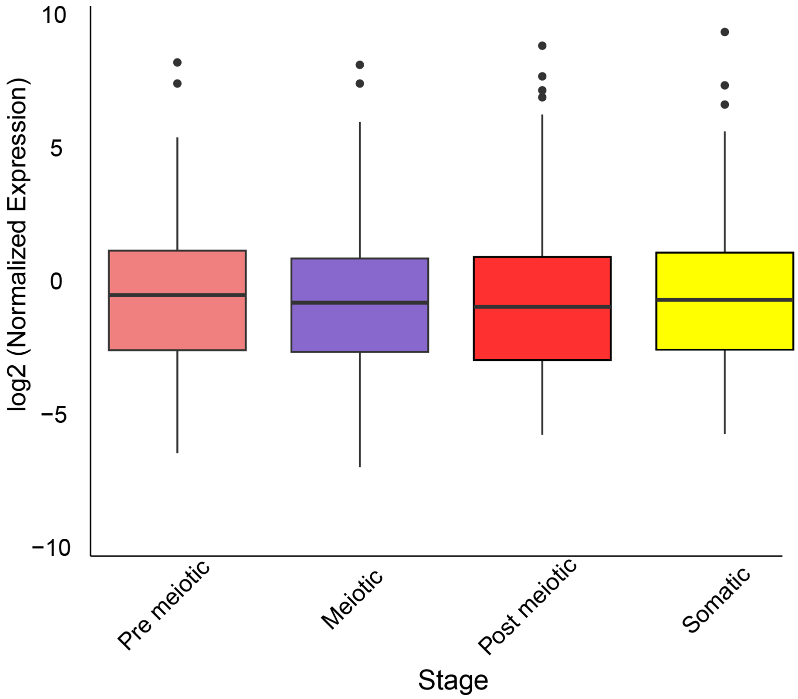


**Supplemental Figure 7. Broadly expressed genes are maintained across all stages of spermatogenesis.** Bulk RNA-seq data from three tissues brain, liver, and testis were compared to identify genes that were expressed testis specifically. When we filtered out 20 testis specific X-linked genes, we still identify similar expression across all stages of spermatogenesis $P > 0.05$ for all comparisons, Kruskal-Wallis and Dunn's test.

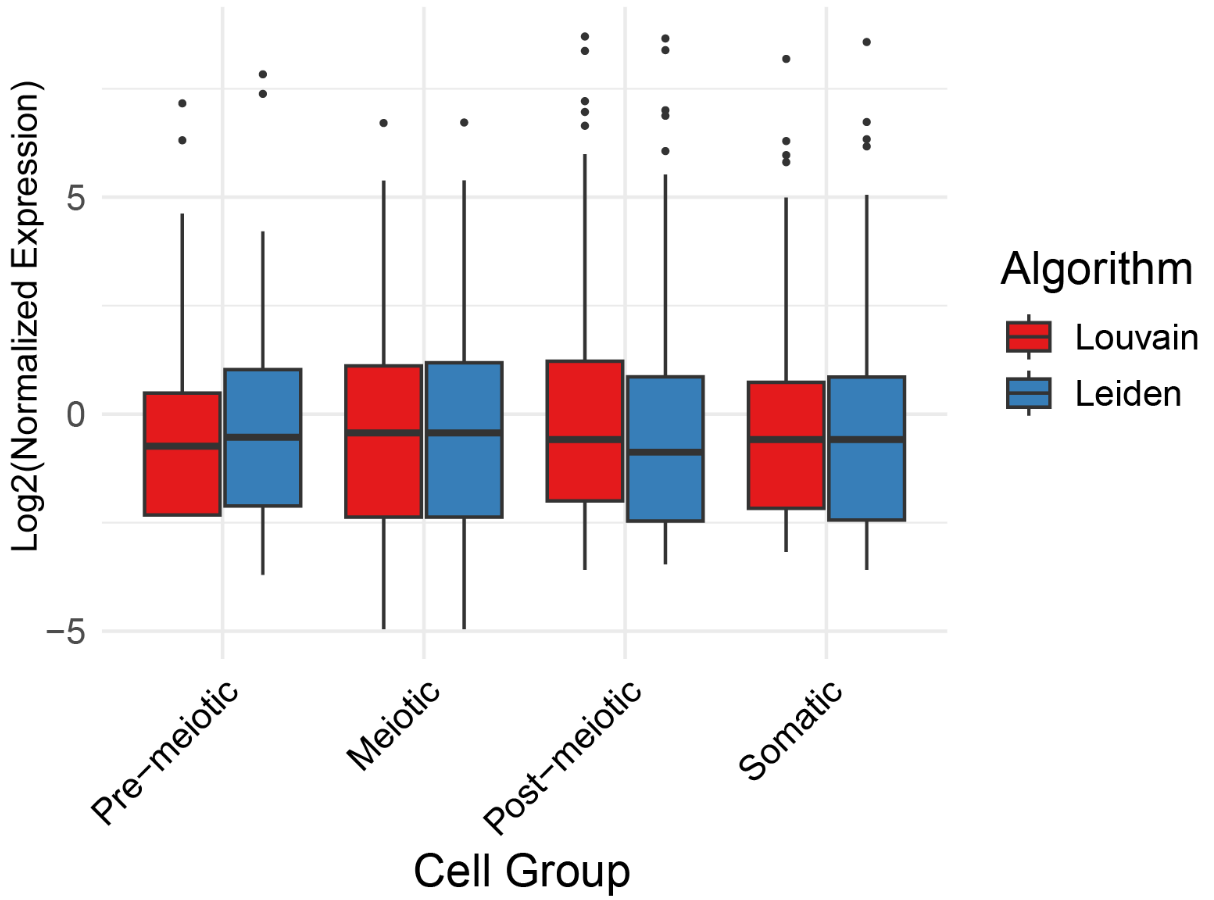
Log2(Normalized Expression)
5
0
−5
Pre−meiotic
Meiotic
Post−meiotic
Somatic
Cell Group
Algorithm
Louvain
Leiden

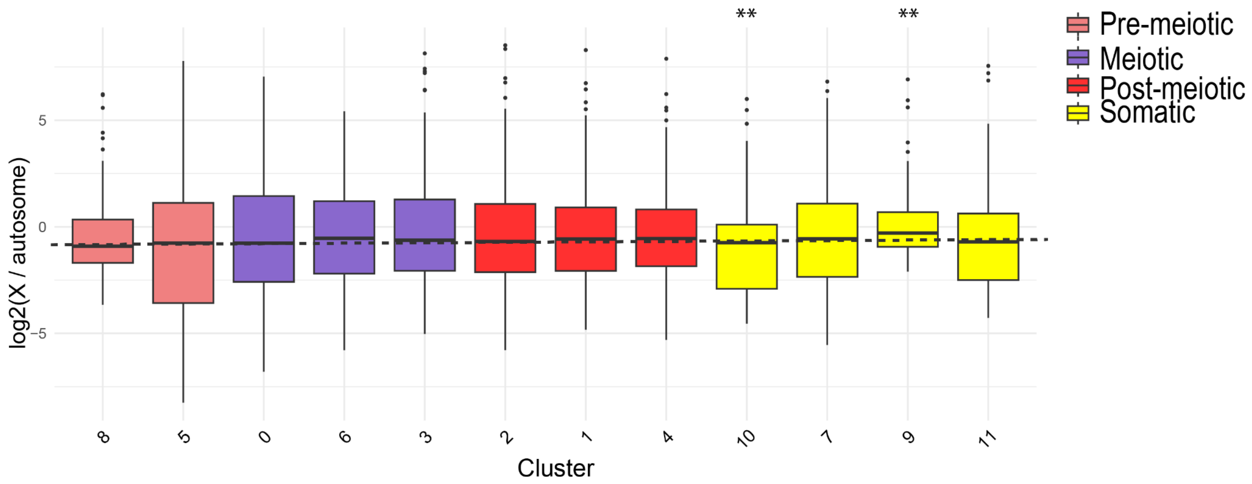
log2(X / autosome)
5
0
-5
**
**
8
5
0
6
3
2
1
4
10
7
9
11
Cluster
Pre-meiotic
Meiotic
Post-meiotic
Somatic

**Supplemental Figure 8. Lack of MSCI is robust across distinct cell clustering algorithms and software.** Top: Normalized X expression was compared between two distinct clustering algorithms in Seurat. X expression was similar across major cell groups for both algorithms (P = 0.441, Kruskal Wallis test. Bottom: Cell clustering was reanalyzed with a separate software package (scanpy) and re-annotated using cell markers. Scanpy clustering was performed using all three samples. Normalized X expression is similar across all stages of spermatogenesis. Only two somatic clusters had variable expression (P < 0.05, Kruskal wallis and Dunn's test).

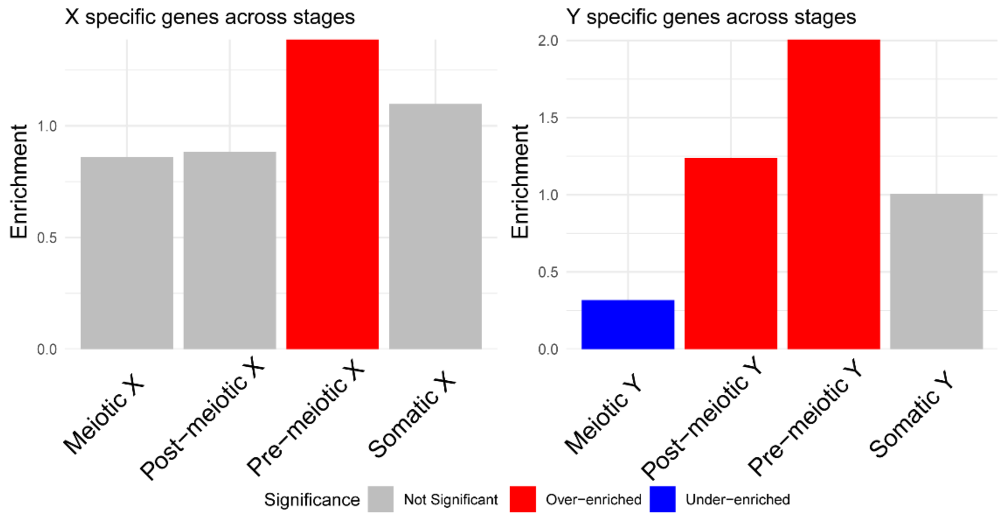


**Supplemental Figure 9. Enrichment of X- and Y- mono-allelic expressed genes across stages of spermatogenesis.** Gametolog specific expression was identified for the 4 main cell types of spermatogenesis by aggregated counts from all clusters. Genes with transcripts aligned to only one sex chromosome per cluster were identified. Mono-allelic expression was confirmed using bulk RNA-seq from three replicates of meiotically active testis. Enrichment of mono-allelic expression across cell stages was calculated using a hypergeometirc test comparing the number of mono-allelic expressed genes observed to expected based on the number of genes expressed in that cell type. X- only expression was overenriched in pre-meiotic cells ( $P < 0.01$). Y-only expression was over-enriched in pre-meiotic and post-meiotic cell types ( $P < 0.01$), and under-enriched in meiotic cell types ($P < 0.01$).

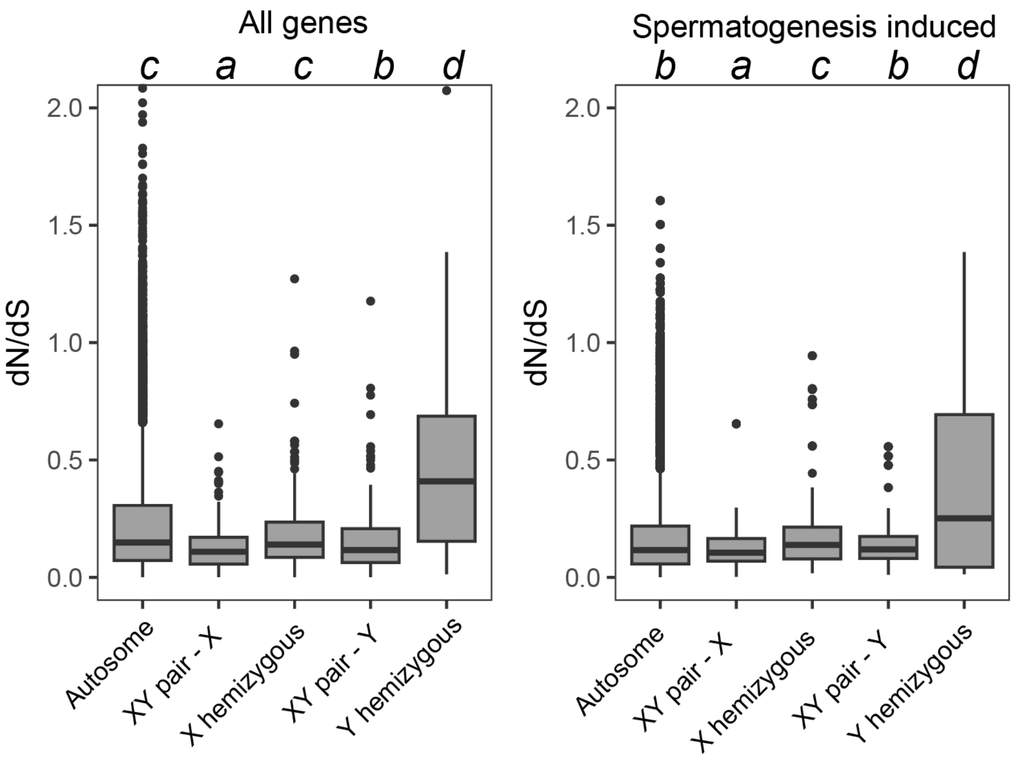


**Supplemental Figure 10.** $d_N/d_S$ **across all induced genes in major cell groups of spermatogenesis.** $d_N/d_S$ was calculated between ninespine and threespine stickleback for genes on autosomes, X hemizygous genes, X and Y gametologs, and Y homozygous genes. scRNA-seq was used to identify genes that were induced in four major cell groups. Letters denote statistical comparisons across all remaining stages and categories based on Kruskal Wallis and Dunn's test. Autosomes have the highest count with 2,262 genes followed by X hemizygous genes with 50 genes, X gametologs with 31 genes, Y gametologs with 31 genes, and Y hemizygous genes with just 4 genes.

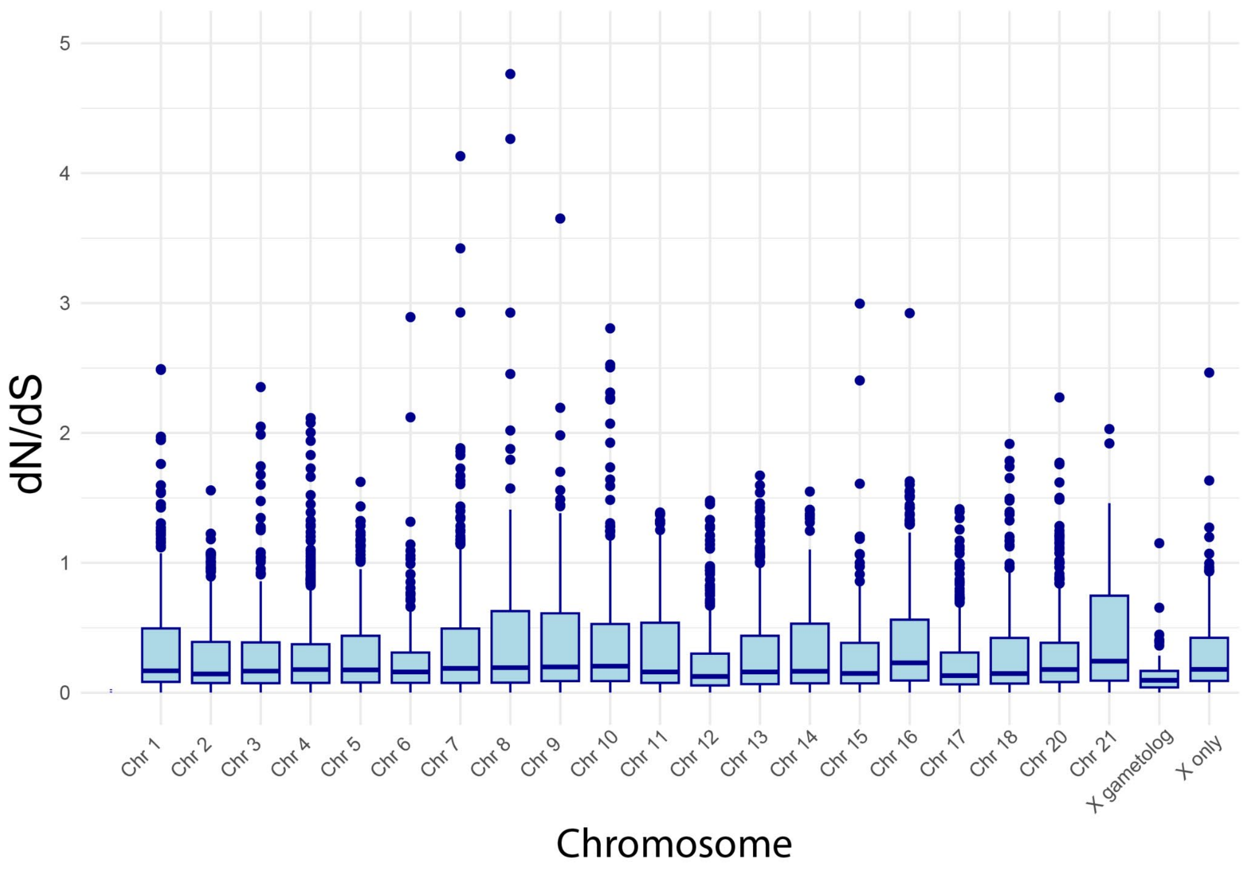


**Supplemental Figure 11.** $d_N/d_S$ **between threespine and ninespine stickleback for all chromosomes.** Both X gametologs and Chr 12 (Ninespine sex chromosomes) have lower dN/dS than median autosomal ($P < 0.01$, Kruskal Wallis and Dunn's test). All other comparisons are not statistically different ($P > 0.05$ for all pairwise comparisons).

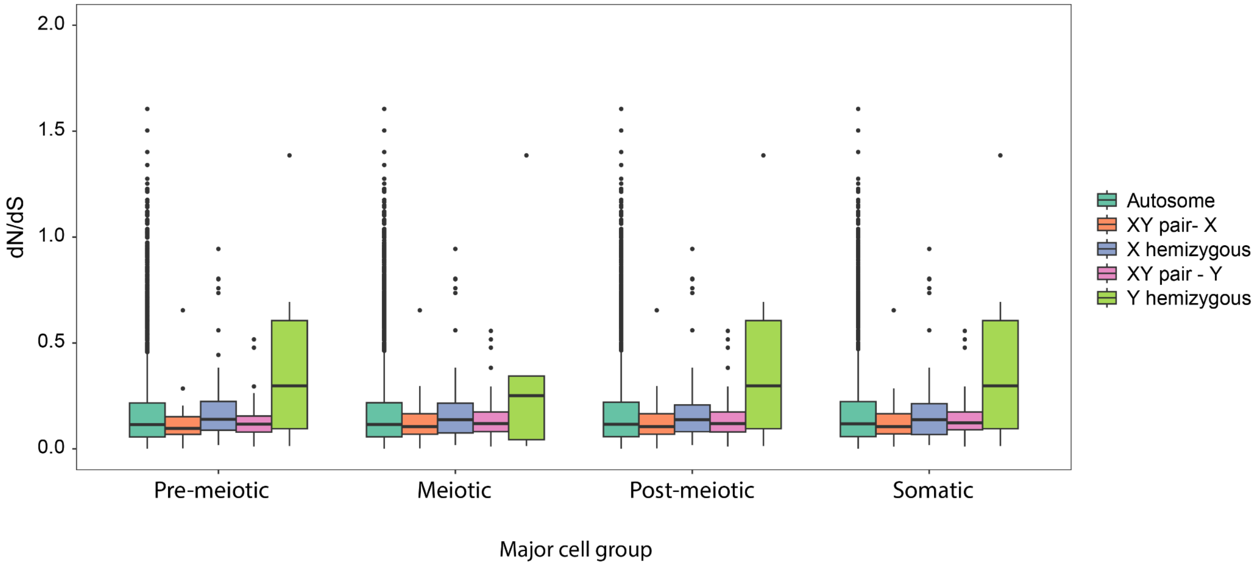


**Supplemental Figure 12. Similar signatures of selection across spermatogenesis.** $d_N/d_S$ was calculated between ninespine and threespine stickleback for genes on autosomes, X hemizygous genes, X and Y gametologs, and Y homozygous genes. scRNA-seq was used to identify 2378 genes that were induced in four major cell groups. There were no significant differences between stages for each chromosome group (P = 0.501 Kruskal Wallis and Dunn's test). The dataset includes genes that were induced and had an alignment between threespine and ninespine stickleback cross four developmental stages for each chromosome category. Autosomes have the largest sample sizes, with 414, 465, 340, and 1,043 genes across the four stages, respectively. X gametologs include 16, 3, 5, and 7 genes; X-only genes have 24, 9, 9, and 8 genes; Y gametologs consist of 16, 3, 5, and 7 genes; and Y-only genes are the smallest group, with just 2 genes in each of the last two stages.

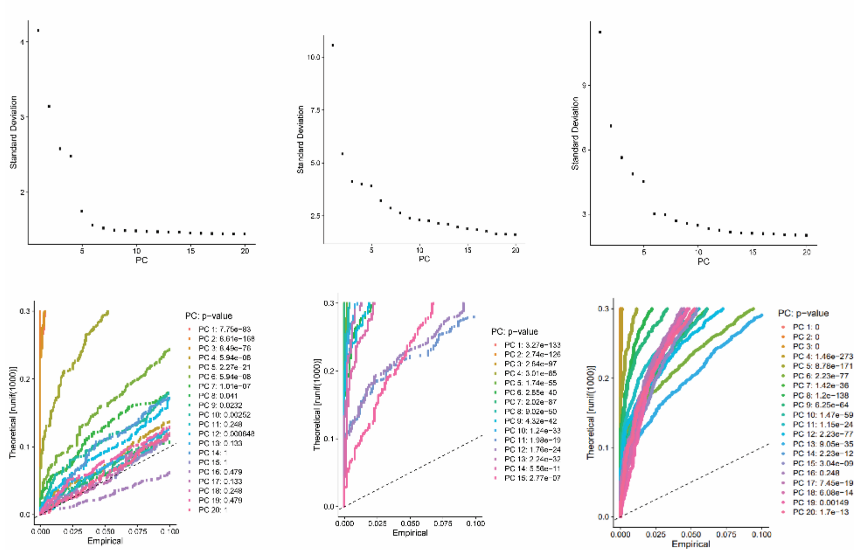


**Supplemental Figure 13. Identification of significant principal components for scRNA-seq**

Elbow plots and jackstraw simulations were generated to assess the number of significant principle components for each sequencing experiment.

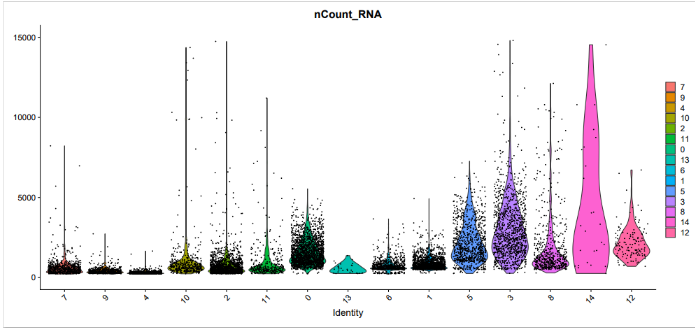


**Supplemental Figure 14. Number of reads per cell type.**

Number of RNA molecules identified per cell within each cell cluster were counted using Seurat.
